# Clinically relevant AAV8-*PEX1* gene therapy preserves retinal integrity and function long-term in a murine model of Zellweger spectrum disorder

**DOI:** 10.64898/2026.05.11.723906

**Authors:** Samy Omri, Erminia Di Pietro, Devin S McDougald, Shaghayegh Zokaei, Jean Bennett, Joseph G Hacia, Nancy E Braverman, Catherine Argyriou

**Affiliations:** Research Institute of the McGill University Health Centre, Montreal, QC, Canada; Center for Advanced Retinal and Ocular Therapeutics, F.M. Kirby Center for Molecular Ophthalmology, Perelman School of Medicine, University of Pennsylvania, Philadelphia, PA, USA; Department of Cancer Biology, Keck School of Medicine, University of Southern California, Los Angeles, CA, USA; Department of Human Genetics, McGill University, Montreal, QC, Canada; Vision Center, Department of Surgery, Children’s Hospital Los Angeles, Los Angeles, CA; The Saban Research Institute, Children’s Hospital Los Angeles, Los Angeles, CA; USC Roski Eye Institute, Department of Ophthalmology, Keck School of Medicine, University of Southern California, Los Angeles, CA

**Keywords:** PEX1, Zellweger spectrum disorder, peroxisome biogenesis disorder, inherited retinal diseases, retinal gene therapy, AAV gene therapy, metabolic gene therapy

## Abstract

Inherited retinal diseases (IRDs) are a heterogeneous group of genetic disorders that cause progressive vision loss. A subset of IRDs is associated with ubiquitously expressed genes involved in fundamental cellular processes, often resulting in multisystem disease. Among these is Zellweger spectrum disorder (ZSD), caused by pathogenic variants in *PEX* genes required for peroxisome biogenesis and function. There are no proven targeted disease-modifying treatments for ZSD, and it is unclear whether localized restoration of peroxisome function is sufficient to mitigate retinal degeneration. We previously demonstrated that *HsPEX1* retinal gene augmentation therapy in a mouse model of mild ZSD homozygous for the murine equivalent (PEX1-p.[Gly844Asp]) of the most common deleterious allele in patients (*PEX1*-c.[2528G>A], PEX1-p.[Gly843Asp]), improved retinal electrophysiological response. Here, we present a comprehensive, dose-range evaluation of a re-designed, clinically relevant AAV8-delivered *HsPEX1* subretinal gene therapy, employing expanded outcome measures. We observed a significant improvement in functional vision, retinal response, photoreceptor structure, retinal pigment epithelium integrity, subretinal inflammation, and peroxisomal metabolites, durable to the endpoint of 6 months post single subretinal injection. These studies provide preclinical proof-of-concept that localized retinal gene replacement can mitigate vision loss in peroxisome-mediated IRD.

## INTRODUCTION

Inherited retinal diseases (IRDs) comprise a heterogeneous group of genetic disorders that cause progressive vision loss, primarily through dysfunction of rod and cone photoreceptors and the retinal pigment epithelium (RPE). Collectively associated with over 260 causal genes, IRDs affect more than 2 million individuals worldwide and are a major cause of inherited blindness ^1, 2^. Because most are driven by loss-of-function (LOF) variants and the retina is readily accessible for local intervention, IRDs have emerged as leading clinical targets for gene augmentation therapies ^3, 4^.

While most IRDs are caused by deleterious variants in genes with retina-specific functions, a subset involves ubiquitously expressed genes that regulate fundamental cellular processes, often resulting in syndromic or multisystem disease ^5^. These diseases present an added therapeutic challenge, as it often remains unclear whether systemic dysfunction contributes to retinal pathology. The peroxisomal biogenesis disorders in the Zellweger spectrum (PBD-ZSD) exemplify this group of IRDs and are an autosomal recessive condition caused by pathogenic LOF variants in *PEX* genes required for peroxisome assembly and matrix protein import ^6, 7^. This impairs critical lipid metabolic pathways, including very long-chain fatty acid (VLCFA) β-oxidation and the synthesis of docosahexaenoic acid (DHA)–containing phospholipids and plasmalogens, as well as multiple other pathways such as reactive oxygen and nitrogen species metabolism ^8–10^. Patients with Zellweger spectrum disorder (ZSD) frequently develop a multisystemic disease with progressive retinal degeneration characterized by early pigmentary changes, macular atrophy, and markedly reduced electroretinographic responses ^7, 11, 12^. No proven disease-modifying therapies are currently available for ZSD.

To study ZSD pathology and test potential therapies, we developed the knock-in *Pex1*^2531G>A/2531G>A^ (PEX1-p.[Gly844Asp];[Gly844Asp]), or PEX1-G844D, mouse model for mild ZSD ^13^. This model recapitulates the common ZSD-causing variant, *PEX1*-c.[2528G>A] (PEX1-p.[Gly843Asp]), and key human disease features including progressive retinopathy ^14, 15^, growth retardation, and liver disease ^16–18^. In an initial proof-of-principle study, we found that adeno-associated virus serotype 8 (AAV8)-mediated retinal *HsPEX1* gene augmentation resulted in a sustained improvement of retinal electrophysiological response ^19^. However, this study used a non-clinically translatable vector design at only a single dose, lacked longitudinal functional vision assessments, and preceded the identification of key pathological features, including dorsal RPE atrophy, progressive photoreceptor loss, and delayed-onset inflammation ^15^, that now represent essential preclinical endpoints. Thus, the extent to which *HsPEX1* retinal gene augmentation can preserve retinal structure, modulate disease-associated inflammation, and achieve long-term therapeutic benefit in this model remains unclear.

Here, we evaluate a clinically relevant AAV8-mediated subretinal gene therapy delivering human *PEX1* cDNA to determine whether localized restoration of peroxisomal function is sufficient to improve visual function, stabilize photoreceptor structure, preserve RPE integrity, modulate retinal inflammation, and correct biochemical hallmarks of peroxisomal dysfunction in the PEX1-G844D mouse model. For biochemical assessments, we incorporated micro dissection of whole neural retina and RPE to limit the possible contamination from surrounding tissues in our initial proof-of-principle study ^19^. In addition, we included *in vitro* and *in vivo* dose-response studies to enable translational development. By integrating functional, structural, and molecular endpoints, we directly test whether retinal degeneration driven by a systemic metabolic disorder can be mitigated through localized retinal gene augmentation. These studies address a pivotal question: whether the effective treatment of vision loss in peroxisome-mediated IRD requires systemic correction, or if retinal-specific rescue can preserve or enhance visual function. More broadly, our work establishes a conceptual framework for treating other multisystemic IRDs and supports the feasibility of targeting the retina in a significant subset of the patient population.

## RESULTS

### Therapeutic vector design

To enable future clinical translation, we re-engineered the therapeutic vector used in our initial proof-of-principle studies ^19^. We maintained the AAV8 serotype for its efficient targeting of photoreceptor and RPE cells in mice ^20, 21^, pigs ^22^, and nonhuman primates ^23, 24^, and incorporated a modified chicken β-actin promoter fused with a CMV enhancer (CBh) and a synthetic polyA recognition sequence (AAV8.*PEX1*, **Figure 1A**, left). This design maximizes expression by maintaining the CMV enhancer while incorporating the more stable chicken β-actin promoter, a combination that has been used clinically and shown efficacy ^25, 26^. Modifications to the polyA recognition sequence result in a shorter construct to accommodate the large (3,852 base pair) codon-optimized human *PEX1* gene, creating a total construct size of 5,074 base pairs ^27, 28^. As a control for AAV8 transduction, we used an identical vector backbone to deliver *GFP* instead of *HsPEX1* (AAV8.*GFP*). Genome integrity was confirmed by long-read sequencing of whole AAV genomes, which demonstrated that 53% of AAV8.*PEX1* capsids and 78% of AAV8.GFP capsids contained the full-length sequence in correct orientation (**Figure 1A**, right).

**Figure 1.**
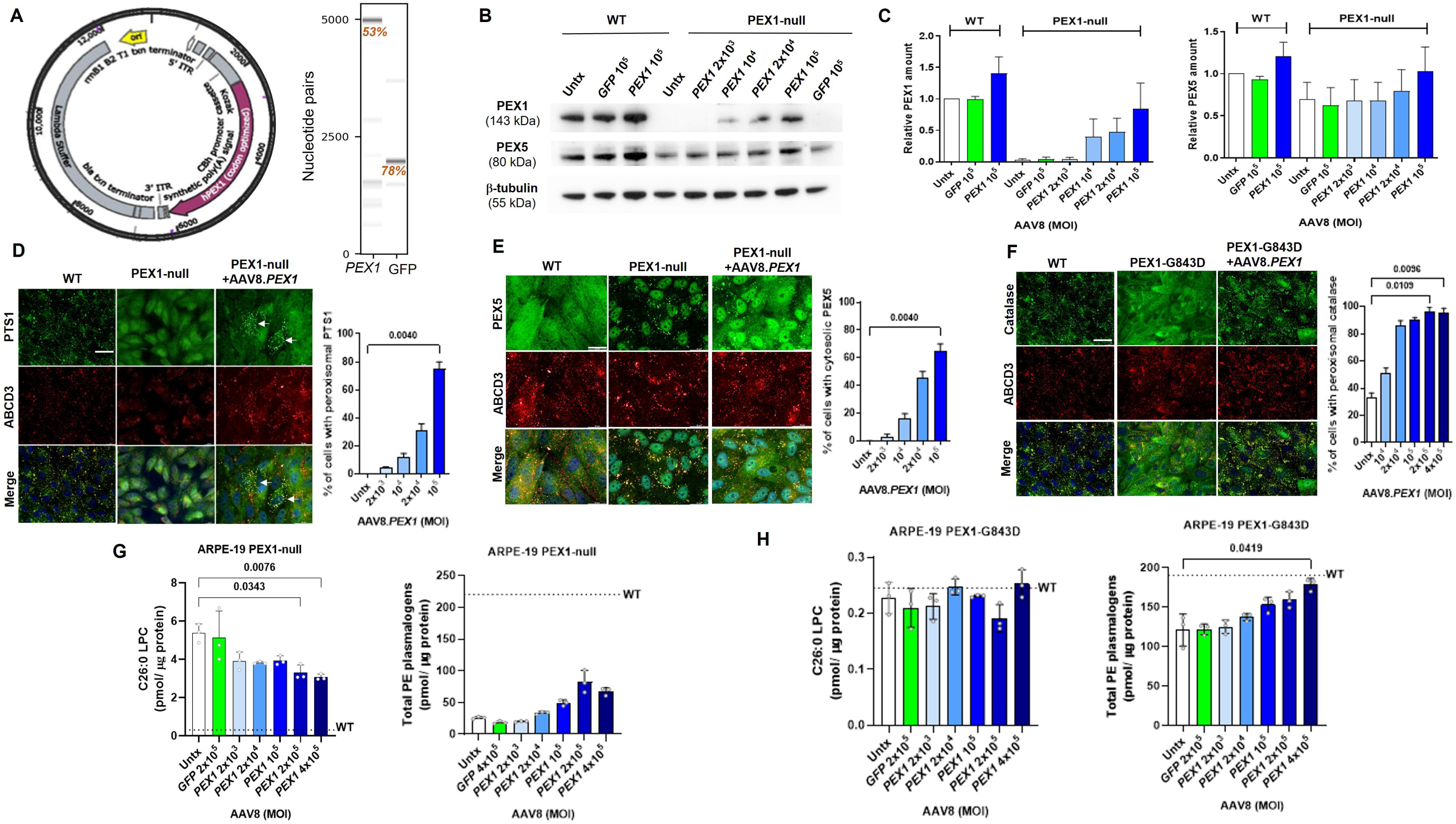
*In vitro* validation of the AAV8.*PEX1* therapeutic vector. **(A)** Left: Schematic of the *PEX1* gene therapy vector featuring the human *PEX1* transgene driven by a chicken β-actin promoter / cytomegalovirus enhancer (CBh) and a synthetic polyadenylation signal within an AAV proviral plasmid backbone. Right: Virtual gel generated based on read length histograms from full-length sequencing of AAV8.*PEX1* and AAV8.GFP vectors. Percent of full-length reads for PEX1 (5,074 nucleotides) and GFP (1,974 nucleotide) constructs are indicated in red. **(B)** Immunoblot analysis of **c**ontrol (WT) and *PEX1***-**null ARPE-19 cell lysates to assess PEX1 and PEX5 protein levels following transduction with AAV8.*PEX1* or AAV8.*GFP.* **(C)** Protein levels quantified by band densitometry and normalized to the β-tubulin loading control; 15ug of protein was loaded per lane. **(D, E)** Immunofluorescence analysis of WT, PEX1-null (untreated), and AAV8.*PEX1*-treated ARPE-19 cells co-labeled for PTS1 and ABCD3 **(D)** or PEX5 and ABCD3 (E), with quantification shown to the right of each panel. White arrows indicate co-localization of PTS1 with the peroxisomal marker ABCD3 in AAV8.*PEX1*-treated cells (phenotypic recovery). **(F)** Immunofluorescence analysis of WT and PEX1-G843D ARPE-19 cells co-labeled for catalase and ABCD3, with quantification shown at right. Images were acquired by fluorescence microscopy at 60x magnification. PTS1, PEX5, or catalase (green) and peroxisome membrane protein ABCD3 (red); colocalization (yellow); DAPI nuclear staining (blue); scale bar, 20µm. n=3 independent experiments with ≥100 cells from ≥6 fields of view quantified per experiment. **(G-H)** LC-MS/MS quantification of peroxisomal metabolites, C26:0-lysophosphatidylcholine (LPC) and phosphatidylethanolamine (PE) plasmalogens, in WT (untreated), **(G)** PEX1-null (untreated and AAV8.*PEX1*-treated) ARPE-19 cells, and **(H)** PEX1-G843D (untreated and AAV8.*PEX1*-treated) ARPE-19 cells, 14 days after transduction. N=3 independent experiments for all; Kruskal–Wallis test with Dunn’s multiple comparison post hoc test; P<0.05 indicated; Mean (SD) shown. Viral dose is expressed as multiplicity of infection (MOI; vg/cell).

### In vitro vector validation and dose-response studies

For *in vitro* validation, we used an immortalized human retinal pigment epithelium line with differentiated properties (ARPE-19) ^29^. This line is amenable to AAV8 transduction and grows in a monolayer, facilitating evaluation by immunofluorescence microscopy (IF). Biallelic *PEX1*-null and *PEX1*-c.[2528G>A] (PEX1-p.[Gly843Asp]) ARPE-19 cell lines were created using CRISPR-Cas-9 gene editing and confirmed by *PEX1* gene sequencing and immunoblot analysis. The PEX1-null line represents a complete loss-of-function system in which any degree of vector-driven PEX1 protein expression or phenotypic rescue is readily detectable, and was thus used to provide maximal dynamic range and assay sensitivity for vector validation. In contrast, the PEX1-G843D ARPE-19 line represents a hypomorphic, partial loss-of-function model designed to recapitulate the residual PEX1 activity present in the majority of *PEX1* patients and the correspondingly milder ZSD phenotype, and, appropriately, show more modest baseline abnormalities. To validate our therapeutic vector, we analyzed PEX1 protein levels, peroxisomal import, and peroxisome metabolite levels following vector transduction in the dose-response studies outlined below. Statistical analyses demonstrating significant dose response trends for all *in vitro* studies are presented in **Table S1**.

#### Protein levels

We used PEX1-null ARPE-19 cells to demonstrate dose-dependent increase in PEX1 protein levels. As the peroxisome matrix protein import receptor PEX5 is degraded when PEX1 is impaired ^30^, we also quantified PEX5 levels following transduction. Cells were transduced at four doses spanning multiplicity of infection (MOI) 2 x 10^3^ to 10^5^ vector genomes (vg) per cell and analyzed by immunoblot after 4 days (**Figure 1B**). In the parent cell line (wild-type cells), a mean 40% increase in PEX1 and 20% increase in PEX5 protein levels was observed at MOI 10^5^ of AAV8.*PEX1*, the only dose tested (**Figure 1C**). In *PEX1*-null cells, PEX1 protein became detectable at MOI 10^4^ and increased dose-dependently from 37% to 84% of wild-type levels. Baseline PEX5 protein levels were 70% of wild-type and increase to wild-type levels was achieved at the highest vector dose. Transduction with AAV8.GFP did not affect PEX1 or PEX5 levels in WT or *PEX1*-null ARPE-19 cells.

#### Peroxisome import/export

We evaluated peroxisome recovery in *PEX1*-null ARPE19 cells transduced with 4 doses of AAV8.*PEX1* and visualized after 6 days by IF. In wild-type control cells, proteins with the C-terminal <u>p</u>eroxisome targeting <u>s</u>ignal 1 (PTS1) motif are localized to peroxisomes, co-localizing with the peroxisomal membrane protein ABCD3. In *PEX1-*deficient cells, PTS1-containing proteins are localized to the cytosol and degraded. AAV8-mediated *PEX1* expression restored PTS1 protein localization in *PEX1*-null ARPE-19 cells in a dose-dependent manner (**Figure 1D**). At the highest dose, PTS1 import was recovered in 75% of cells. As a complementary measure, we assessed localization of the PEX5 shuttle protein. In wild-type cells, PEX5 is primarily cytosolic. In *PEX1*-deficient cells, PEX5 appears punctate and co-localizes with the peroxisome membrane protein ABCD3, indicating it is trapped at the peroxisome membrane and not recycled for additional rounds of import. Following AAV8.*PEX1* transduction, PEX5 localization was restored to the cytosol in a dose-dependent manner, with 65% of cells recovered at the highest dose (**Figure 1E**). Recovery of PEX5 localization provides direct evidence of restored PEX1 function, which is to remove PEX5 from the peroxisome membrane. In wild-type cells, transduction did not alter PTS1 or PEX5 localization, which remained correct (cytosolic) in all cells. We were unable to assess the effects of AAV8.GFP transduction in IF analyses, as GFP expression interferes with visualization.

To model milder ZSD phenotypes, we used ARPE-19 cells homozygous for the *PEX1*-c.[2528G>A] variant (PEX1-G843D). During characterization, we observed that this line does not exhibit a robust defect in PTS1 protein or PEX5 localization. We thus used catalase localization, which is typically perturbed by the mildest peroxisome import dysfunction due to its non-canonical PTS1 motif ^31^. At baseline, catalase was localized to peroxisomes in 33% of PEX1-G843D ARPE-19 cells. As the PEX1-G843D ARPE-19 cells proliferated more rapidly than the *PEX1*-null cells (diluting therapeutic transgene-expressing cells with each division), we expanded the vector doses to include 5 doses spanning MOI 10^4^ to 2 x 10^5^ to increase the likelihood of achieving phenotypic recovery over baseline. We omitted the 2 x 10^3^ vg dose, as no PEX1 protein increase or phenotypic improvement was detectable at this dose in *PEX1*-null ARPE-19 cells. In PEX1-G843D ARPE-19 cells, AAV8.*PEX1* transduction corrected catalase localization in a dose-dependent manner, with over 90% of cells recovered at MOI 2 x 10^5^ (**Figure 1F**). There was no observable effect of transduction in wild-type cells. The highest dose tested (MOI 4×10 ) caused moderate toxicity.

#### Downstream peroxisome function

To confirm downstream recovery of peroxisome function, we evaluated peroxisome metabolite levels routinely used for clinical diagnosis by liquid chromatography-tandem mass spectrometry (LC-MS/MS) ^32^. In general, impaired peroxisomal β-oxidation results in increased C26:0 very long chain fatty-acid (C26:0-VLCFA) levels. In addition, peroxisomes are required for the synthesis of plasmalogens, a class of ether phospholipids ^33^. VLCFAs (represented by C26:0 lysophosphatidylcholine, C26:0-LPC) and phosphatidylethanolamine (PE) plasmalogen levels were measured by LC-MS/MS 14 days post AAV8.*PEX1* transduction in control, *PEX1-*null, and PEX1-G843D ARPE-19 cells. At baseline, C26:0-LPC levels were elevated over 17-fold in the *PEX1*-null cells compared to wild-type. These levels were partially recovered at all doses, with a 40% reduction occurring at the highest doses (**Figure 1G, left**). Total PE plasmalogens were reduced to 12% of wild-type levels in *PEX1*-null cells at baseline. A dose-dependent trend toward partial recovery was observed following therapeutic vector transduction, with the two highest doses resulting in a 2.6 – 3.2 fold increase in PE plasmalogen levels (**Figure 1G, right**). Transduction with AAV8.*GFP* did not affect the lipid levels measured (only the lowest and highest doses tested). In PEX1-G843D ARPE-19 cells, baseline C26:0-LPC levels were not elevated compared to wild-type, and AAV8 transduction had no effect (**Figure 1H, left**). Total PE plasmalogens were reduced by 37% at baseline in PEX1-G843D ARPE-19 cells, and were increased at the highest dose, at which they no longer differed from WT levels (**Figure 1H, right**).

#### Murine cell rescue

To confirm that our therapeutic vector recovers peroxisome function in a *Pex1*-deficient murine background, we used primary retinal pigment epithelium (RPE) cells cultured from PEX1-G844D mice and WT littermate controls. At baseline, 5% of PEX1-G844D cells exhibited cytosolic PEX5 localization (**Figure 2A**). Transduction with AAV8.*PEX1* recovered PEX5 localization in a dose-dependent manner, with 80% of cells exhibiting cytosolic PEX5 localization at the maximum dose (MOI 10^5^) without apparent toxicity (**Figure 2A**). MOI 2 x 10^5^ recovered PEX5 localization in all surviving cells but resulted in moderate toxicity and thus was not pursued in experimental replicates. Viral transduction did not affect PEX5 localization in WT cells, which remained cytosolic in all cells.

**Figure 2.**
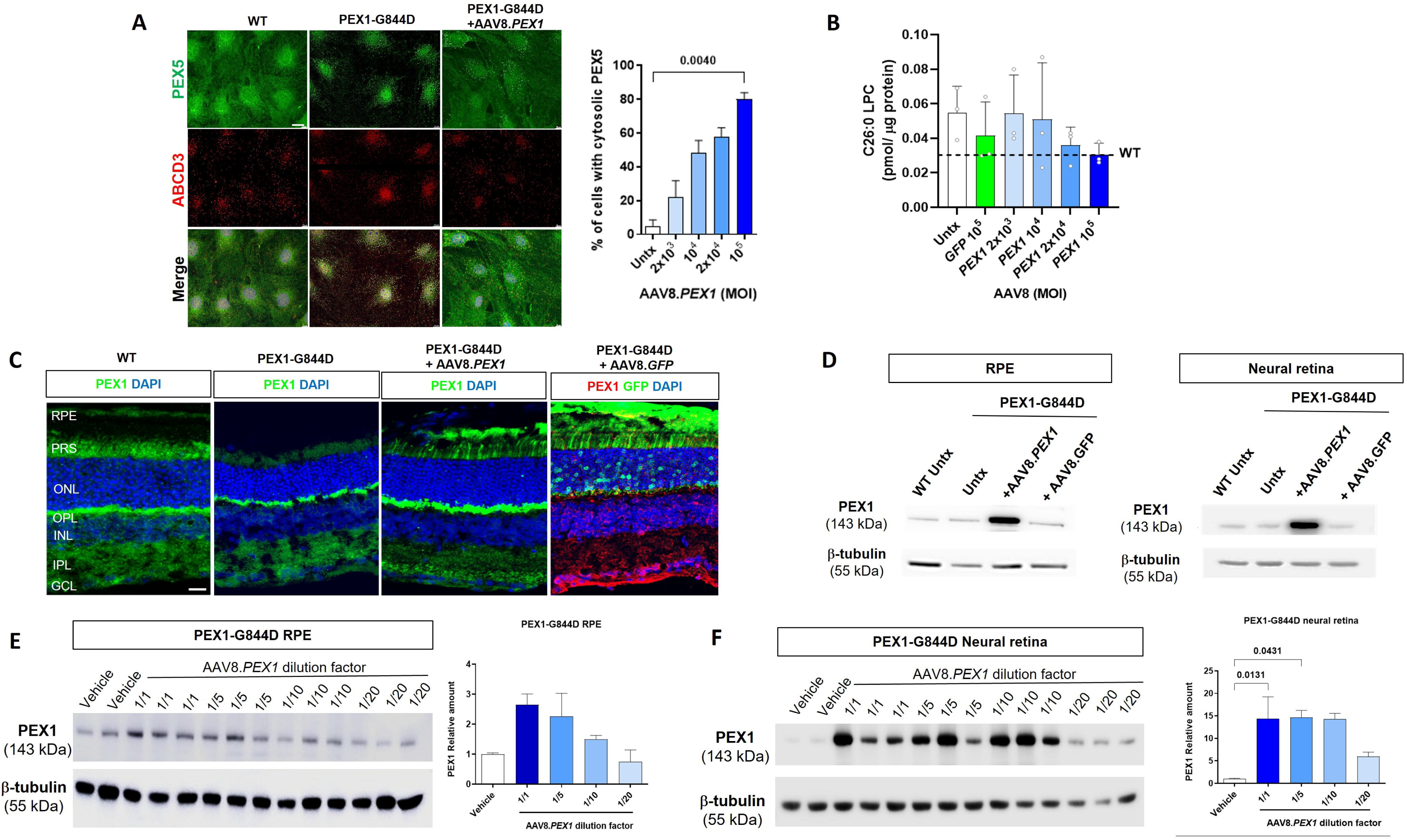
*In vitro* and *in vivo* validation of the AAV8.*PEX1* therapeutic vector in murine tissues. **(A)** Immunofluorescence analysis of WT and PEX1-G844D primary mouse RPE cells treated with various doses of AAV8.*PEX1*, co-labelled for PEX5 (green), ABCD3 (red), and DAPI (blue), with quantification shown at right. Images were acquired by fluorescence microscopy at 60x magnification; Scale bar, 20µm. N=3 independent experiments with ≥100 cells from ≥ 6 fields of view quantified per experiment. Viral dose is expressed as multiplicity of infection (MOI; vg/cell). **(B)** LC-MS/MS quantification of the peroxisomal metabolite C26:0-lysophosphatidylcholine (LPC) in PEX1-G844D primary RPE cells (untreated and AAV8.*GFP-* or AAV8.*PEX1*-treated), 14 days post-transduction. The untreated WT average is shown for reference. **(C)** Representative confocal z-stack images of WT and PEX1-G844D retinal cryosections 2 months after a single subretinal injection of 6.2 x 10^9^ vg AAV8.*PEX1* or AAV8.*GFP*, immunolabelled with PEX1 (green or red) and DAPI (blue); GFP signal is shown in green. PRS, Photoreceptor segments; ONL, outer nuclear layer; INL, inner nuclear layer; GCL, ganglion cell layer. Scale bar, 20 µm. Data are presented as mean and SD, n = 3 mice per group. **(D)** Immunoblot analysis of whole neural retina or whole RPE lysates to assess PEX1 protein levels 2 months post single subretinal injection of 6.2 x 10^9^ vg AAV8.*PEX1* or AAV8.*GFP.* **(E, F)** Immunoblot analysis of PEX1-G844D whole neural retina (**E**) or whole RPE **(F)** lysates to assess PEX1 protein levels 2 month after a single subretinal injection of serial dilutions of AAV8.*PEX1*. Doses tested (vg/eye): 6.2 x 10^9^ (1/1), 1.24 x 10^9^ (1/5), 6.2 x 10^8^ (1/10), 3.1 x 10^8^ (1/20), and 6.2 x 10^7^ (1/100). Protein levels were quantified by band densitometry and normalized to the β-tubulin loading control; 15ug of protein was loaded per lane. N=3 independent experiments; Statistical analysis was performed using a Kruskal–Wallis test with Dunn’s multiple-comparisons post hoc test; P < 0.05 indicated. Data are presented as mean (SD).

To assess downstream recovery of peroxisome function, we evaluated common clinical biomarkers. In PEX1-G844D primary RPE cells, C26:0-LPC levels trended toward elevation at baseline (1.8-fold) compared to cultured cells from WT littermate controls. However, this difference was not statistically significant. Following AAV8.*PEX1* transduction at the highest dose (MOI 10^5^), C26:0-LPC levels trended toward reduction (**Figure 2B**). Total PE plasmalogen levels were not reduced at baseline in PEX1-G844D primary RPE cells and were not used as an outcome.

### In vivo vector validation

#### Protein levels

To confirm vector expression and localization following subretinal delivery, we visualized PEX1 and GFP in mouse retinal sections by immunohistochemistry 4 weeks post injection in mice treated at 1 month of age. For both therapeutic and control vectors, 6.2 x10^9^ vg were administered, corresponding to 1µL of undiluted AAV8.*PEX1*. At baseline, endogenous MmPEX1 immunofluorescence was reduced in PEX1-G844D mice compared to WT. Subretinal AAV8.*PEX1* delivery increased total PEX1 protein (MmPEX1 + HsPEX1) within the injected region, including the RPE and photoreceptor inner and outer segments, mimicking the endogenous MmPEX1 localization pattern observed in WT mice (**Figure 2C**). In eyes injected with AAV8.*GFP*, GFP was expressed in photoreceptor outer segments (OS), outer nuclear layer (ONL), and the outer plexiform layer (OPL), as expected with subretinal AAV8-mediated delivery (**Figure 2C, S1A**). Immunoblotting of micro dissected whole RPE and neural retina demonstrated increased PEX1 protein levels compared to untreated, vehicle-injected, and AAV8.*GFP*-injected tissues (**Figure 2D**). To assess retinal coverage, we examined GFP distribution in RPE flatmounts from PEX1-G844D mice injected with AAV8.*GFP*. We observed consistent GFP expression in the dorsal pole, and variable expression throughout the remaining retina (**Figure S1A**). GFP expression was still present 5 months post injection in both neural retinal sections and RPE flatmounts, despite progressive morphological changes and cell loss in PEX1-G844D retinas (**Figure S1B, S1C**).

To determine whether vector dose correlates with increased PEX1 protein levels *in vivo*, we administered AAV8.*PEX1* at the undiluted dose (1/1) plus 1/5, 1/10, and 1/20 dilutions (6.2 x 10^9^ vg, 1.24 x 10^9^ vg, 6.2 x 10^8^ vg, and 3.1 x 10^8^ vg, respectively), all in 1µL subretinal injection. In RPE, a dose-dependent relationship was observed, with the highest dose producing the greatest PEX1 band density (∼2.7-fold increase in PEX1 protein levels), and the lowest dose showing no increase (**Figure 2E**). In neural retina, the three higher doses produced a 14-fold increase in PEX1 protein levels, and the lowest dose a 6-fold increase (**Figure 2F**). Statistical analyses demonstrating significant dose-response trends are presented in **Table S1**. These observations are consistent with known AAV8 tropism; both RPE and photoreceptors were transduced and expressed the therapeutic protein, with higher transduction efficiency in the neural retina ^34^.

### In vivo dose optimization

To establish the optimal vector dose for expanded preclinical efficacy studies, we tested the effects of various doses on functional and structural recovery in PEX1-G844D mice. We administered AAV8.*PEX1* by subretinal injection at the undiluted dose of 6.2 x 10^9^ vg per eye (1/1), as well as 1.24 x 10^9^ vg, 6.2 x 10^8^ vg, and 6.2 x 10^7^ vg per eye, corresponding to 1/5, 1/10, and 1/100 dilutions, respectively. No intervention (Untx) and 1 µL vehicle injection served as negative controls. Mice were injected at 1 month of age and evaluated 2 months later. Statistical analyses demonstrating significant dose-response trends for all *in vivo* studies are presented in **Table S1**.

#### Functional vision

The Striatech OptoDrum platform was used to assess functional vision following vector administration using optomotor reflex testing (OMR). Under ambient light (photopic) conditions, full-contrast spatial acuity was 25% lower in untreated PEX1-G844D mice compared to WT littermate controls, and the highest vector dose improved spatial acuity by 34% (**Figure 3A**). Treatment did not alter scotopic spatial acuity, which remained 30% lower than WT. Contrast sensitivity, however, was improved 2-fold at the 1/1 and 1/5 doses, reaching WT levels. Neither high-dose vector administration nor injection significantly affected functional vision measures in WT, except for reduced contrast sensitivity in the vehicle-injected group (**Figure S2A**). Detailed statistical analyses of these observations are presented in **Table S2** and **Table S3**.

**Figure 3.**
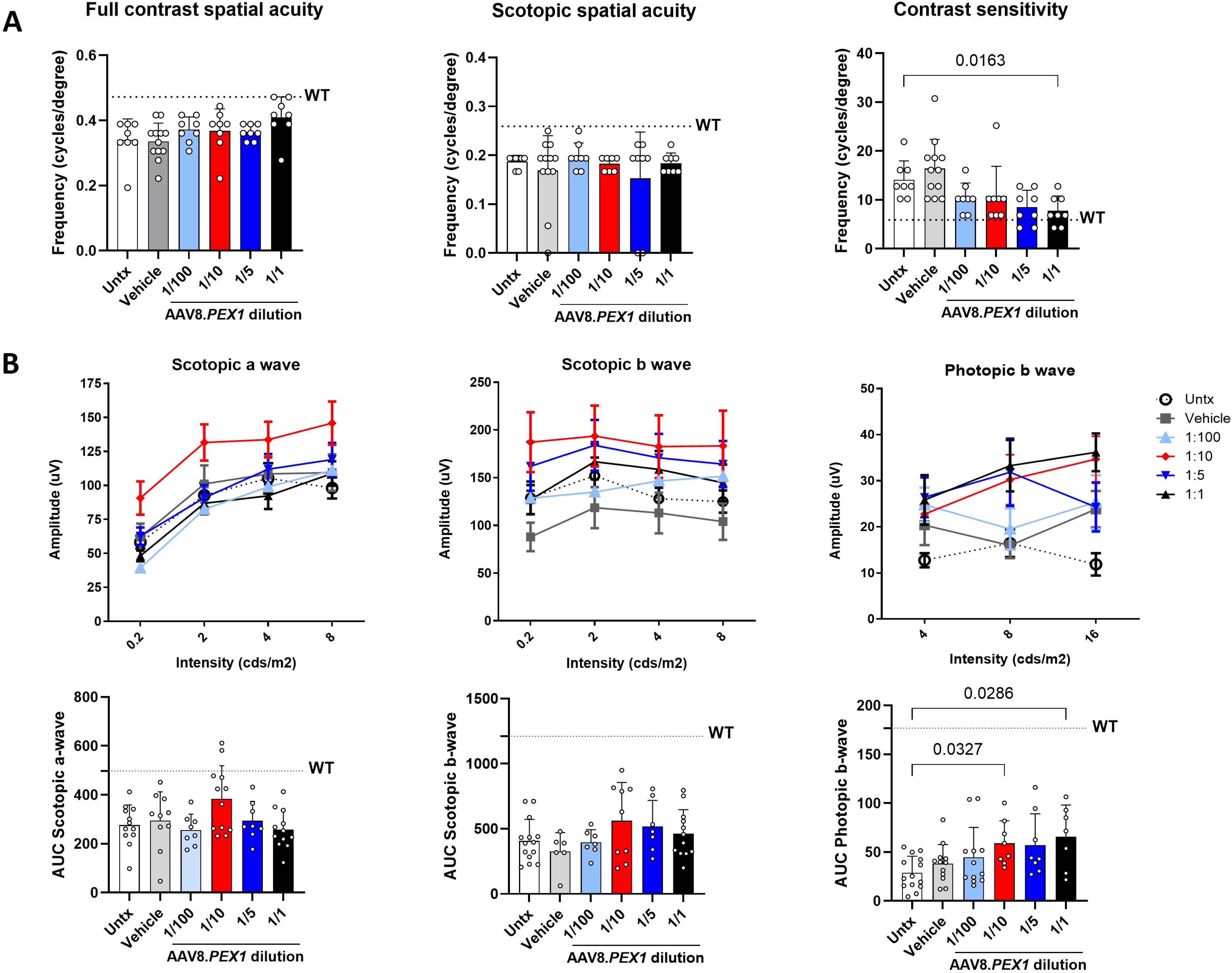
Dose-dependent effects of AAV8.*PEX1* on retinal function in PEX1-G844D mice. AAV8.*PEX1* was administered by subretinal injection to 1-month-old PEX1-G844D mice, and assessments were performed 2 months post-treatment (age 3 months). Doses tested (vg/eye): 6.2 x 10^7^ (1/100), 6.2 x 10^8^ (1/10), 1.24 x 10^9^ (1/5), and 6.2 x 10^9^ (1/1). Average values from untreated WT littermates are shown as a reference (dotted line). **(A)** Optomotor reflex testing of functional vision, including full contrast spatial acuity under ambient and scotopic conditions (reported as the highest spatial frequency perceived) and contrast sensitivity under ambient light (reported as the lowest contrast perceived). **(B)** Full-field flash electroretinography (ffERG) responses were quantified as amplitudes of scotopic a-wave, scotopic b-wave, and photopic b-wave at increasing stimulus intensities. The area under the curve (AUC) for each recording is shown below each plot. Data represent 4–8 mice per group (8–12 eyes). Statistical analysis was performed using a Kruskal–Wallis test with Dunn’s multiple-comparisons post hoc test; P < 0.05 indicated; Data are presented as mean (SD).

#### Retinal response

The Diagnosys Celeris platform was used to assess retinal responses by full-field flash electroretinography (ffERG) following vector administration. In PEX1-G844D mice, there was a statistically significant improvement in scotopic a-wave at the 1/10 dose, as well as photopic b-wave at the 1/1 and 1/10 doses. Overall, the three higher doses presented a trend toward ffERG improvement, and there was no statistically significant effect of vehicle injection (**Figure 3B**). Detailed statistical analyses of these observations are presented in **Table S2**.

#### Photoreceptor structure

Labelling of cone OS using peanut agglutinin (PNA) was reduced in untreated PEX1-G844D mutants compared to WT littermates. Representative images suggest increased PNA labeling following AAV8.*PEX1* treatment, particularly at the 1/5 and 1/1 doses (**Figure 4A**). Similarly, ONL thickness was reduced by about 15% in untreated PEX1-G844D mice compared to WT (**Figure 4B, 4C, 4D**). The 1/5, 1/10, and 1/100 doses all showed a trend toward improvement, and which was statistically significant at the 1/5 dose, at which ONL thickness was preserved at WT levels (**Figure 4C, 4D**). Treatment had no effect on WT photoreceptor morphology (**Figure S2B, S2C**). Detailed statistical analyses of these observations are presented in **Table S3** and **Table S4**.

**Figure 4.**
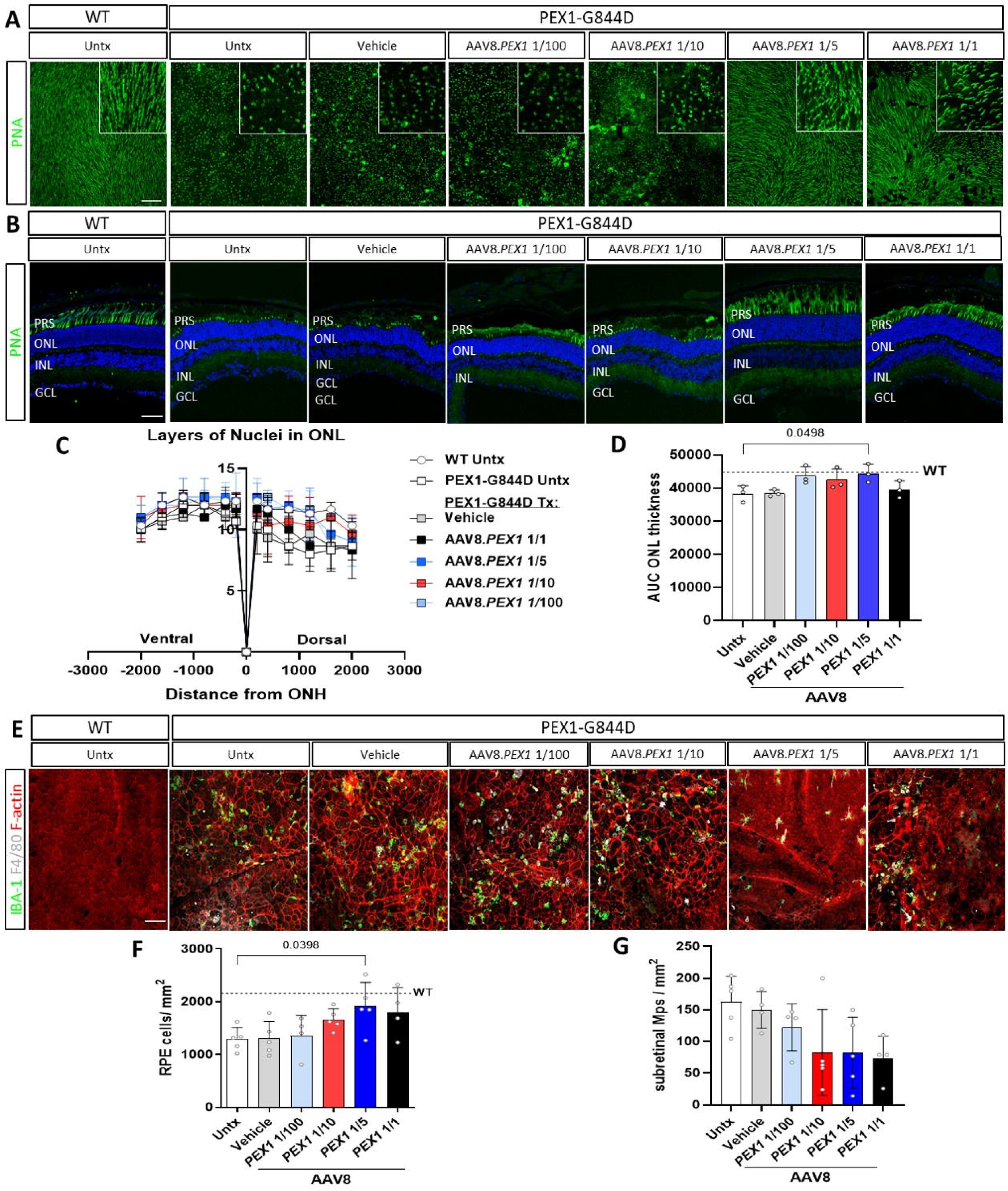
Dose-dependent effects of AAV8.*PEX1* on retinal structure in PEX1-G844D mice. AAV8.*PEX1* was administered by subretinal injection to 1-month-old PEX1-G844D mice, and assessments were performed 2 months post-treatment (age 3 months). Doses tested (vg/eye) were: 6.2 x 10^7^ (1/100), 6.2 x 10^8^ (1/10), 1.24 x 10^9^ (1/5), and 6.2 x 10^9^ (1/1). **(A, B)** Representative confocal z-stack images from untreated littermate controls (WT) and PEX1-G844D mice across treatment groups. **(A)** Neural retina flatmounts (photoreceptor side up) stained with peanut agglutinin (PNA), a lectin that labels the extracellular matrix surrounding cone photoreceptors. Scale bar, 100 µm. **(B)** Retinal cryosections from the same groups stained with peanut agglutinin (PNA) to visualize cone-associated extracellular matrix across retinal layers. Scale bar, 20 µm. **(C)** Spider plot representation of ONL thickness across the entire retinal circumference. **(D)** Area under the curve (AUC) analysis of ONL thickness. Data are presented as mean (SD); n = 3 mice per group. PRS, Photoreceptor segments; ONL, outer nuclear layer; INL, inner nuclear layer; GCL, ganglion cell layer. **(E)** Representative immunofluorescence images of RPE flatmounts (apical side up) from untreated littermate controls (WT), and PEX1-G844D mice across treatment groups. Subretinal immune cells are labeled with IBA1 (green) and F4/80 (white), and RPE morphology is visualized with TRITC-phalloidin staining of F-actin (red). Scale bar, 100 µm. **(F)** Quantification of RPE cell density (cells/mm²). **(G)** Quantification of subretinal immune cell density (IBA1 /F4/80 cells/mm²). n = 4–5 mice per group; Statistical analysis was performed using a Kruskal–Wallis test with Dunn’s multiple-comparisons post hoc test; P < 0.05 indicated. Data are presented as mean (SD).

#### RPE structure and inflammatory features

Untreated PEX1-G844D mice exhibited a 35% reduction in RPE cell density compared to WT, concurrent with subretinal mononuclear phagocyte (macrophage) infiltration (**Figure 4E-G**). AAV8.*PEX1* preserved RPE cell density, with a statistically significant effect at the 1/5 dose (13% lower than WT), and a trend toward improvement at the 1/1 and 1/10 doses (**Figure 4E, 4F**). Similarly, AAV8.*PEX1* produced a significant dose-dependent trend toward reduced subretinal macrophages, although values were variable (**Figure 4G**). At the 1/5 dose, microglia displayed a more ramified morphology, indicating lower inflammatory profile compared to the amoeboid morphology associated with a pro-inflammatory state ^35, 36^ (**Figure 4E**). In WT mice, there was a mild but statistically significant increase in RPE cell density following AAV8.*PEX1* injection (**Figure S2D, S2E**). In addition, AAV8.*PEX1* treatment produced a very low level of subretinal macrophage infiltration (4-13 macrophages per mm^2^) (**Figure S2F, S2G**). Detailed statistical analyses of these observations are presented in **Table S3** and **Table S4**.

### Expanded long-term preclinical efficacy study of AAV8.PEX1 in PEX1-G844D mice

Our next step was to perform an expanded, long-term preclinical efficacy assessment of AAV8.*PEX1* in a large mouse cohort. Based on our *in vivo* dose optimization studies, we selected the 1/5 dose (1.24 x 10^9^ vg) as it produced the most consistent retinal preservation across functional and structural outcomes. All mice underwent baseline ffERG assessment at 4 weeks of age and received bilateral subretinal AAV8.*PEX1* injection at 5 weeks of age. No treatment, vehicle injection, and 1.2 x 10^9^ vg AAV8.*GFP* were used as controls, and both WT and PEX1-G844D mice were included in all groups. ffERG and OMR testing were performed at 2, 4, and 6 months post-gene delivery. The experimental endpoint was 6 months post injection (7 months of age), at which point retinal structure and peroxisomal metabolites were analyzed in neural retina and RPE.

### Long-term improvement of functional vision and retinal response post AAV8.PEX1 treatment

#### Functional vision

OMR testing was used to assess spatial acuity and contrast sensitivity under both ambient light and scotopic conditions. Longitudinal analyses showed that improved functional vision was evident at 2 months post treatment in the AAV8.*PEX1*-treated PEX1-G844D mice, and was maintained or improved over 6 months (**Figure S3A**). There was no effect of treatment in WT mice (**Figure S3A**). At the experimental endpoint (7 months of age), spatial acuity in untreated PEX1-G844D mice was >60% below the WT average under both lighting conditions. In PEX1-G844D mice treated with AAV8.*PEX1*, spatial acuity under both lighting conditions improved more than twofold, reducing the disparity with WT to 25% (**Figure 5A**). Moreover, contrast sensitivity under ambient light improved by over threefold, reaching WT levels (**Figure 5B**). Finally, scotopic contrast sensitivity improved by 60% with treatment, approaching WT levels (**Figure 5B**). Detailed statistical analyses of these observations are presented in **Table S5**.

**Figure 5.**
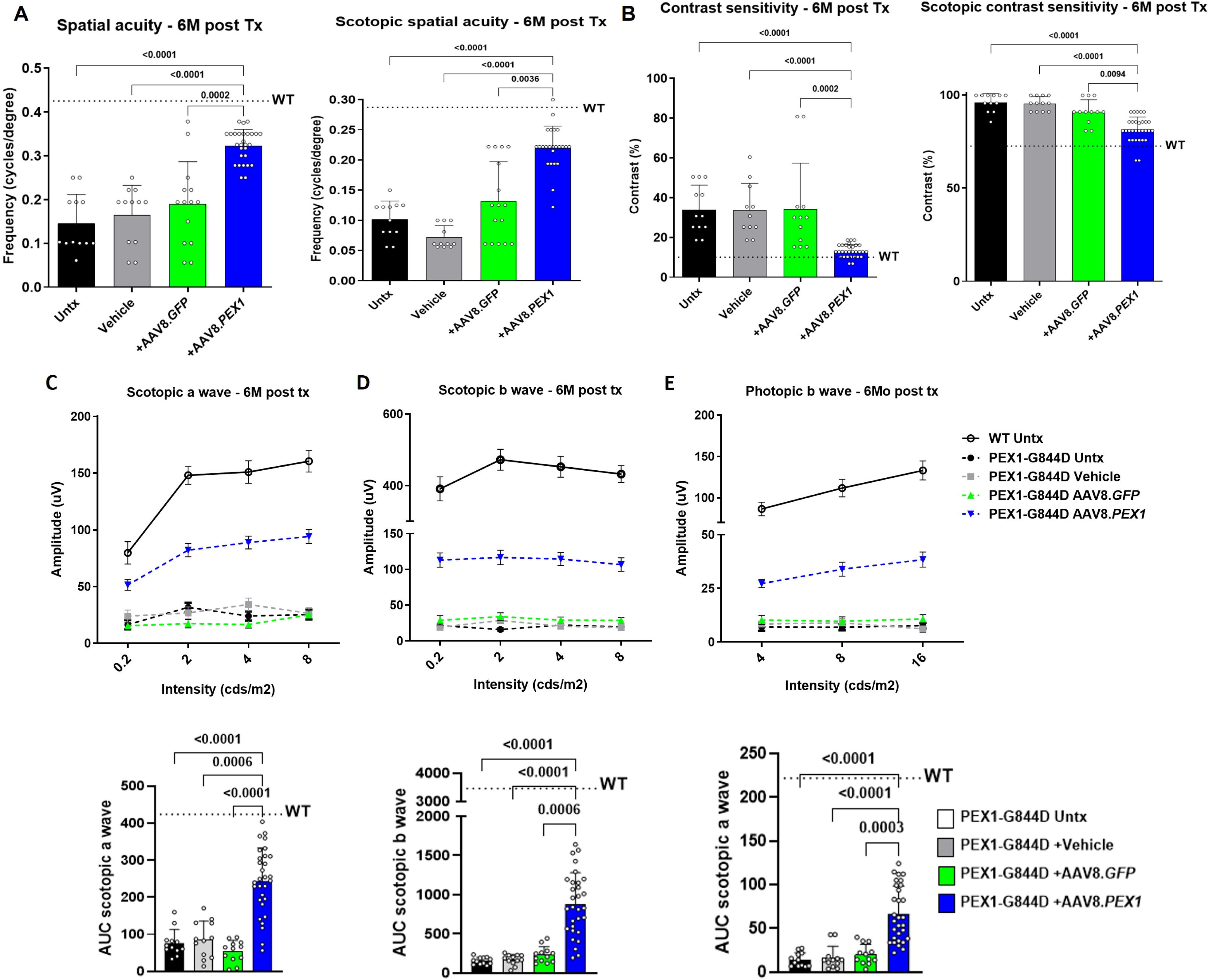
Retinal function 6 months following AAV8.*PEX1* treatment in PEX1-G844D mice. 1.24 x 10^9^ vg/eye AAV8.*PEX1* or AAV8.*GFP* was administered by subretinal injection to 5-week-old PEX1-G844D mice, and assessments were performed 6 months post-treatment (age 7 months). Optomotor reflex testing of functional vision included: **(A)** full contrast spatial acuity under ambient and scotopic conditions (reported at the highest spatial frequency perceived) and **(B)** contrast sensitivity at ambient light and scotopic conditions (reported as the lowest contrast perceived). **(C-E)** Full-field flash electroretinography (ffERG) responses were quantified as amplitudes of **(C)** scotopic a-wave, **(D)**, scotopic b-wave, and **(E)** photopic b-wave at increasing stimulus intensities. Average values from untreated WT littermates are shown as a reference (dotted line). The corresponding area under the curve (AUC) is shown below each plot. Data represent 6–15 mice per group (12–30 eyes). Statistical analysis was performed using a Kruskal–Wallis test with Dunn’s multiple-comparisons post hoc test; P < 0.05 indicated. Data are presented as mean (SD).

#### Retinal response

ffERG was used to assess retinal response to light stimuli. Qualitatively, at 2 months post injection, scotopic b-wave and photopic b-wave responses were over 2-fold higher in AAV8.*PEX1*-treated PEX1-G844D mice compared to all other treatment groups. At 4 months post injection, all ffERG measures were 3-to 5-fold greater in AAV8.*PEX1*-treated PEX1-G844D mice, and this effect was sustained through 6 months (**Figure S3B**). Photopic ffERG in AAV8.*PEX1*-treated PEX1-G844D mice improved 2-to 2.4-fold between baseline and endpoint. Formal statistical testing was performed at study endpoint. For scotopic ffERG, the decline over 6 months was reduced by more than half in AAV8.*PEX1*-treated PEX1-G844D mice. ffERG responses declined moderately over time in wild-type mice across all treatment groups, consistent with age-related decline (**Figure S3B**). At the experimental endpoint, AAV8.*PEX1* recovered retinal response to 30-40% of wild-type levels in PEX1-G844D mice across scotopic a-wave (**Figure 5C**), scotopic b-wave (**Figure 5D**), and photopic b-wave responses (**Figure 5E**), representing an average response more than 5-fold greater than untreated PEX1-G844D mice. Detailed statistical analyses of these observations are presented in **Table S5**.

### Long-term photoreceptor preservation and reduced gliosis post AAV8.PEX1 treatment

In PEX1-G844D mice treated with therapeutic vector, ONL thickness and cone OS length were significantly preserved at the experimental endpoint. We assessed photoreceptor layer integrity by scoring ONL thickness across the entire retinal length (**Figure 6A**), observing complete preservation in the ventral retina, and mild decline on the dorsal side in AAV8.*PEX1*-treated PEX1-G844D mice (**Figure 6B**). The average ONL thickness in treated PEX1-G844D mice was not statistically different from WT, whereas ONL thickness was reduced by 75% in all other treatment groups. Treatment had no effect of intervention on ONL in WT retinas (**Figure S4A**). In addition, AAV8.*PEX1* improved cone-associated matrix labelling, indicating recovered OS formation (**Figure 6C**). Finally, Müller cell dedifferentiation, measured by glial fibrillary acidic protein (GFAP), a marker of gliosis, was observed in PEX1-G844D retinas not treated with AAV8.*PEX1.* In contrast, GFAP distribution in in AAV8.*PEX1*-treated PEX1-G844D mice resembled WT (**Figure 6D**). Detailed statistical analyses of these observations are presented in **Table S6**.

**Figure 6.**
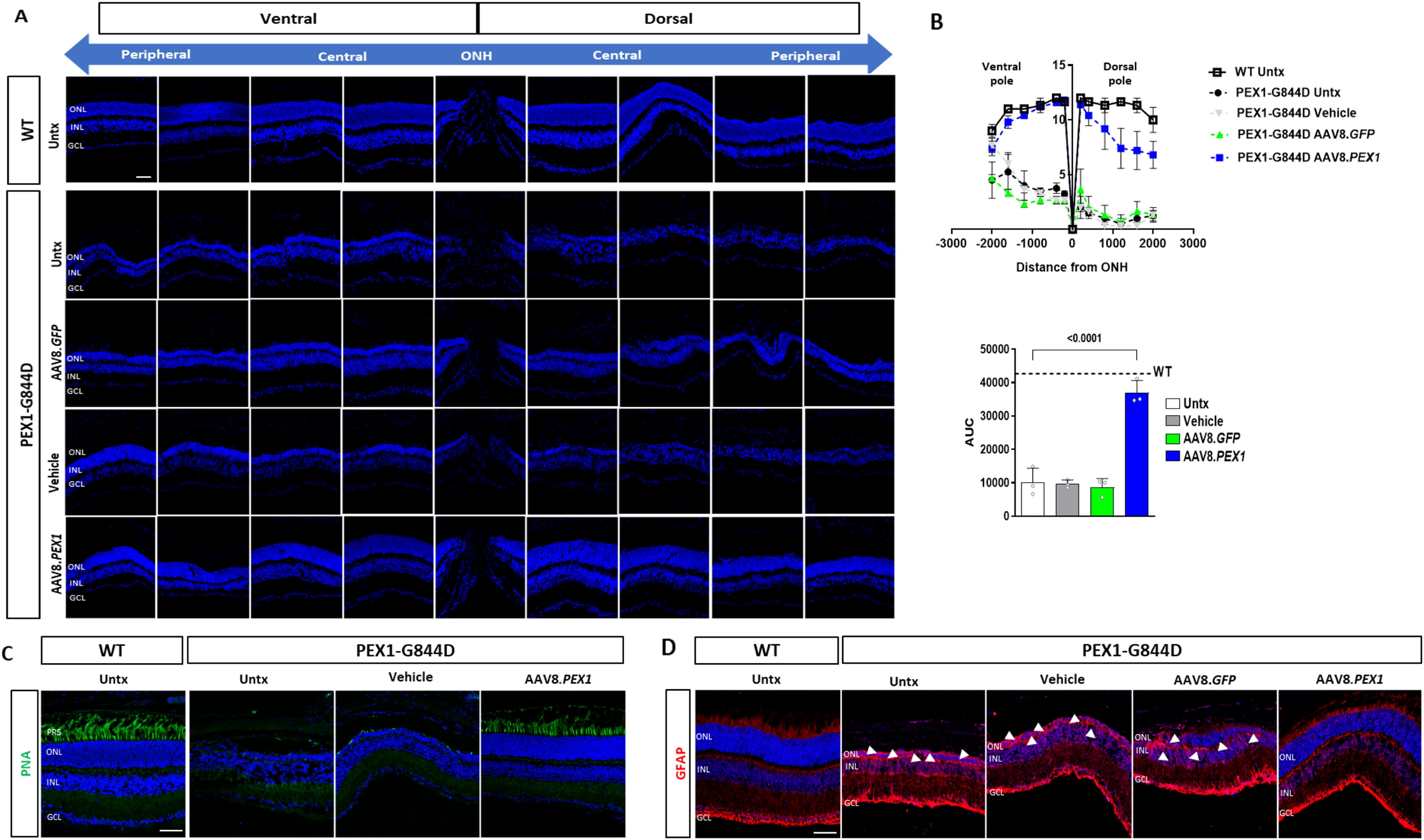
Photoreceptor integrity and gliosis 6 months following AAV8.*PEX1* treatment in PEX1-G844D mice. AAV8.*PEX1* (1.24 x 10^9^ vg/eye) or AAV8.GFP was administered by subretinal injection to 5-week-old PEX1-G844D mice, and assessments were performed 6 months post**-**treatment (age 7 months). **(A)** DAPI-stained retinal cryosections spanning both sides of the optic nerve head (ONH), from peripheral ventral to peripheral dorsal retina, illustrating regional variation in ONL thickness. **(B)** Quantification of ONL thickness across the retinal circumference displayed as a spider plot (n = 3-5 mice per group). The dorsal pole, the most affected region in untreated mutant mice, remained moderately thinned in treated eyes, whereas ventral ONL was better preserved. The area under curve (AUC) is shown below. Statistical analysis was performed using a one way ANOVA followed by Dunnett’s multiple comparisons post hoc test; P < 0.05 indicated. Data are presented as mean ± SD. **(C)** Representative retinal cryosections stained with peanut agglutinin (PNA) to label cone outer segments. **(D)** Representative retinal cryosections labeled for glial fibrillary acidic protein (GFAP, red). Robust Müller glial activation was observed in all mutant groups except AAV8.*PEX1*-treated eyes, which showed reduced GFAP immunoreactivity. Scale bar, 20µm. PRS, Photoreceptor segments; ONL, outer nuclear layer; INL, inner nuclear layer; GCL, ganglion cell layer.

To observe the long-term effects of higher dose AAV8.*PEX1* and AAV8.GFP on ONL integrity, we analyzed outer nuclear layer thickness across the entire retinal length in WT and PEX1-G844D mice 6 months following injection with the undiluted dose of 6.2 x 10^9^ vg/eye. In WT mice, we observed a non-statistically significant reduction in ONL thickness at the injected (dorsal) pole following AAV8 administration, regardless of transgene (**Figure S5A, S5B**). In contrast, PEX1-G844D mice exhibited a trend toward improved ONL thickness with high dose AAV8.*PEX1* administration, particularly in the ventral pole (**Figure S5A, S5C**).

### Long-term effects of AAV8.PEX1 on RPE structure and inflammation

In PEX1-G844D mice treated with therapeutic vector, RPE integrity was significantly preserved at the experimental endpoint (**Figure 7A, 7B**). At 7 months of age, PEX1-G844D RPE exhibits geographic atrophy, characterized by loss of RPE cells leading to patch holes in the RPE layer. To account for potential bias when scoring these regions (i.e. zones lacking tissue will not contain macrophages), we adapted our scoring approach to correct for surface area. We quantified RPE integrity at the experimental endpoint by calculating RPE cell density per surface area (**Figure 7C**). RPE from AAV8.*PEX1*-treated PEX1-G844D mice treated had twofold higher cell density than other mutant treatment group, and only 25% fewer cells than WT. AAV8.*PEX1*-treated PEX1-G844D tissues were almost completely protected from geographic atrophy (**Figure 7D**). We assessed subretinal inflammation by quantifying subretinal mononuclear phagocytes normalized to intact surface area. In AAV8.*PEX1*-treated PEX1-G844D mice, this measure was variable, ranging from no reduction to a 90% reduction compared to other treatment groups (**Figure 7E**). RPE cell density was unaffected in WT mice treated with AAV8.*PEX1*, and two of three retinas analyzed displayed modest subretinal macrophage infiltration (**Figure S4B**). Detailed statistical analyses of these observations are presented in **Table S6**.

**Figure 7.**
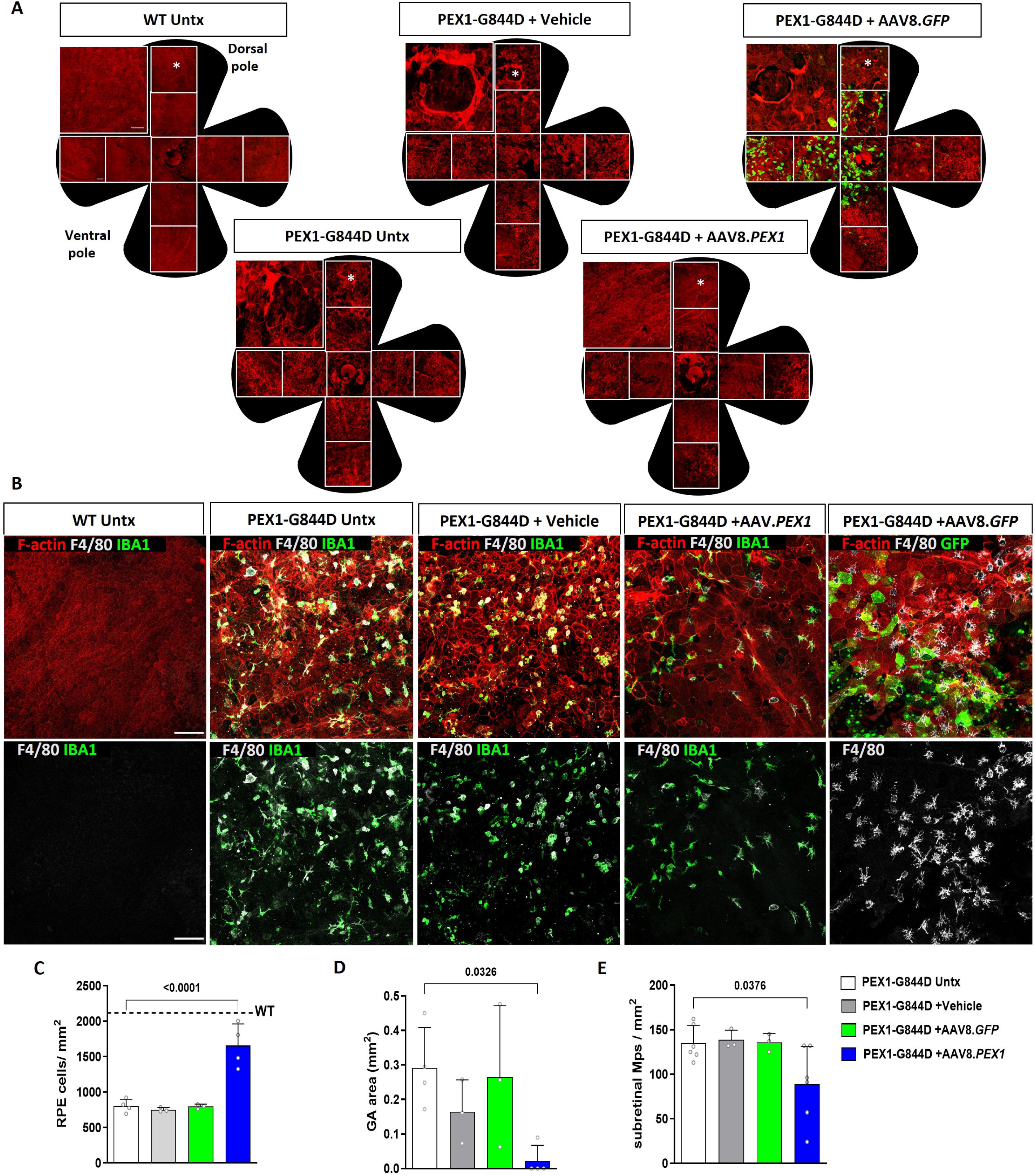
RPE structural integrity and immune cell infiltration 6 months following AAV8.*PEX1* treatment in PEX1-G844D mice. AAV8.*PEX1* (1.24 x 10^9^ vg/eye) or AAV8.*GFP* was administered by subretinal injection to 5-week-old PEX1-G844D mice, and assessments performed 6 months post treatment (age 7 months). Representative confocal z-stack immunofluorescence images are shown. **(A)** RPE flatmounts (four-petal preparations) stained with TRITC-phalloidin (red) to visualize F-actin. GFP fluorescence (green) is present only in the AAV8.*GFP* group, indicating sustained transgene expression. Insets highlight the dorsal pole (asterisk), where focal RPE loss appears as holes consistent with geographic atrophy. **(B)** Representative confocal z-stack images from each group showing triple staining of F-actin (red), IBA1 or GFP (green), and F4/80 (white) in the top row, and corresponding IBA1/F4/80 dual staining in the bottom row to visualize subretinal mononuclear phagocyte infiltration. **(C)** Quantification of RPE cell density (cells/mm²) n = 3–4 mice per group, 6-8 eyes. **(D)** Quantification of geographic atrophy (GA) area (mm²;n = 3–4 mice per group). Data were analyzed using one way ANOVA followed by Dunnett’s multiple comparisons post hoc test. **(E)** Quantification of subretinal mononuclear phagocyte density (cells/mm²) n=3–6 mice per group (6–12 eyes); Scale bar, 100 μm. Statistical analysis was performed using a Kruskal–Wallis test with Dunn’s multiple-comparisons post hoc test; P < 0.05 indicated. Data are presented as mean (SD).

### Long-term effects of AAV8.PEX1 on retinal lipid composition

At the experimental endpoint, neural retina and RPE tissues were isolated for LC-MS/MS analyses. In PEX1-G844D mice, treatment with AAV8.*PEX1* reduced C24:0-LPC and C26:0-LPC levels by over 60% in neural retina compared to other treatment groups (**Figure 8A**). In RPE, C26:0-LPCs were comparably reduced in AAV8.*PEX1*-treated *PEX1*-G844D mice. Relative to neural retina, RPE C24:0-LPC levels were more mildly elevated in untreated PEX1-G844D mice compared to WT. AAV8.*PEX1* treatment produced a trend toward reduction to a level no longer significantly different from WT (**Figure 8B**). Total plasmalogen or phosphoethanolamine (PE) plasmalogen levels were not reduced in PEX1-G844D neural retina or RPE compared to WT, and instead trended toward mild elevation in the neural retina. AAV8.*PEX1* caused a trend toward mild reduction in the PEX1-G844D neural retina compared to untreated, aligning values with WT **(Figure 8A**). PE plasmalogens containing docosahexaenoic acid (DHA, C22:6) were comparable to WT levels in neural retina but decreased by over 50% in PEX1-G844D neural retina and RPE, and were not recovered by AAV8.*PEX1* treatment (**Figure 8A, 8B**). More detailed analyses of PE and phosphatidylcholine (PC) plasmalogen subgroups (**Figure S6**) revealed that AAV8.*PEX1* treatment significantly reduced levels of the pro-inflammatory lipid arachidonic acid (C20:4) in neural retina (**Figure S6A**). Treatment did not affect lipid composition in WT tissues (**Figure 8, Figure S6**). Detailed statistical analyses of biochemical metabolite data are presented in **Table S7**.

**Figure 8.**
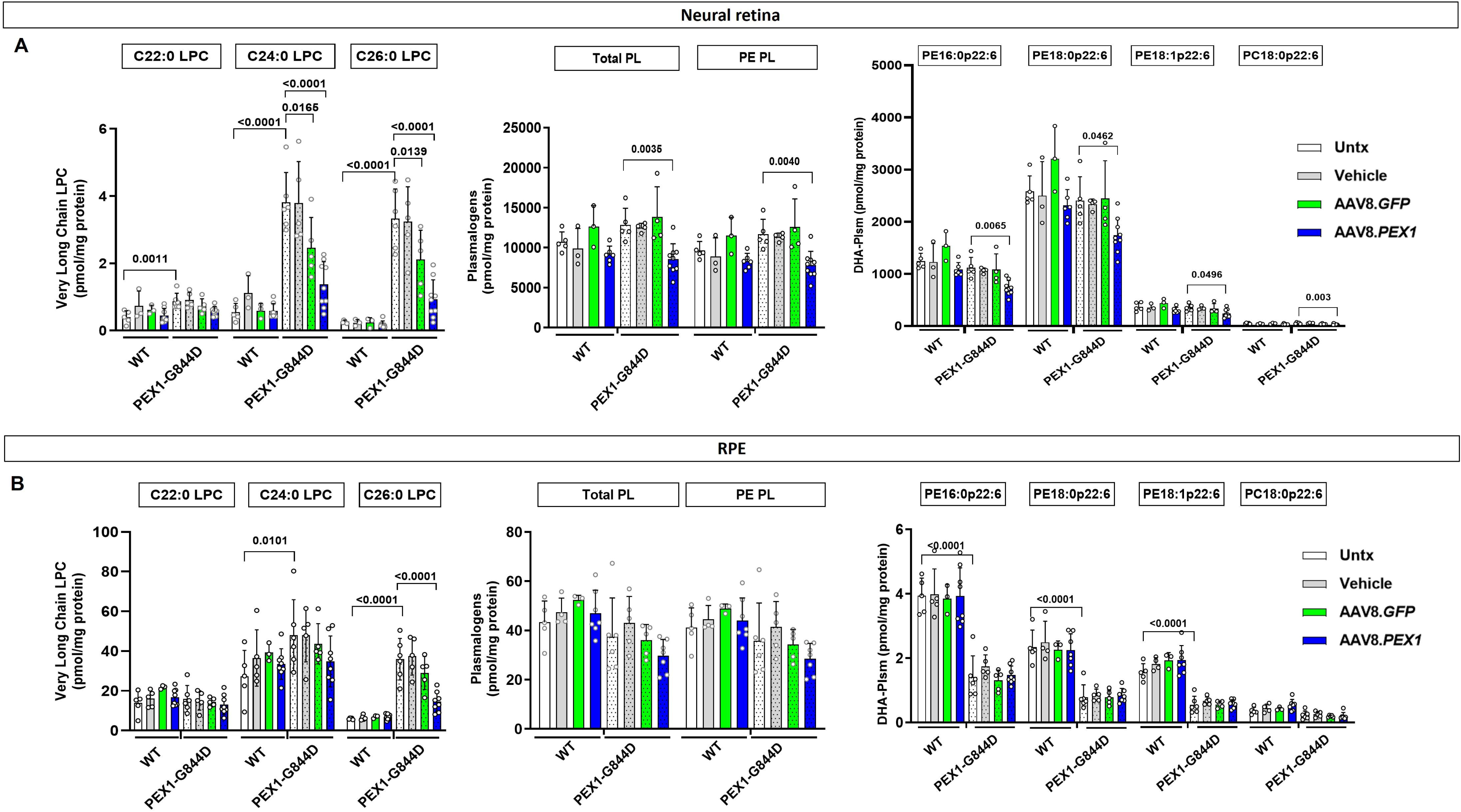
LC-MS/MS quantification of peroxisomal metabolites 6 months following AAV8.*PEX1* treatment in WT and PEX1-G844D mice. Levels of total C26:0-lysophosphatidylcholine (LPC), plasmalogens, and subclasses of phosphatidylethanolamine (PE) plasmalogens, were measured in **(A)** whole neural retina and **(B)** whole RPE from WT and PEX1-G844D mice 6 months after subretinal injection with vehicle, AAV8.*PEX1* (1.24 x 10^9^ vg/eye) or AAV8.*GFP* (age 7 months); N= 5-7 tissues per group. Data were analyzed using two-way ANOVA followed by Tukey’s multiple-comparisons post hoc test (cell-wise); P < 0.05 indicated. Data are presented as mean (SD).

## DISCUSSION

The clinical success of voretigene neparvovec (Luxturna^TM^) has transformed the therapeutic landscape for IRDs primarily confined to the eye, demonstrating that targeted retinal gene augmentation can improve vision, particularly when administered prior to irreversible structural damage ^3^, ^37^. In contrast, IRDs caused by pathogenic variants in ubiquitously expressed genes that affect multiple organ systems (often called syndromic retinal diseases) present a distinct therapeutic challenge, as it often remains unclear whether systemic dysfunction contributes to retinal pathology ^38–40^. ZSD caused by mild *PEX1* deficiency exemplifies this class. Biallelic hypomorphic alleles in genes encoding the peroxisomal exportomer complex (*PEX1*, *PEX6*, and *PEX26*) disrupt peroxisomal metabolic pathways across tissues and produce a complex multisystem phenotype characterized by progressive retinal degeneration and early vision loss, as well as hearing loss, amelogenesis imperfecta, and depending on the level of residual activity, adrenal, liver, and central nervous system involvement ^11, 41–44^.

Here, we demonstrate that localized AAV8-mediated *PEX1* gene augmentation partially restores peroxisomal function in the PEX1-G844D murine retina and yields sustained structural and functional benefits. Importantly, these effects are achieved despite persistent systemic metabolic abnormalities, providing evidence that retinal degeneration in this context is, at least in part, retina-autonomous and amenable to local correction. These findings establish that targeted retinal gene therapy can modify disease progression in a multisystem peroxisomal disorder.

For disorders like ZSD, in which systemic correction remains challenging, the ability to preserve vision through localized intervention represents a clinically meaningful and tractable therapeutic strategy. This is particularly relevant given patient-centered studies identifying vision preservation as a major priority for affected individuals ^45^. As an initial target, *PEX1* is relevant for a large proportion of the global ZSD population, particularly in the intermediate to milder end of the disease spectrum where intervention is most likely to change disease trajectory ^46^. More broadly, our results support a conceptual framework in which tissue-specific gene augmentation can provide therapeutic benefit in a subset of multisystem IRDs, even when the underlying metabolic defect is not corrected systemically.

Compared to our earlier proof-of-concept study in the same homozygous PEX1-G844D model, which used a non-clinical HA-tagged vector and lacked dose evaluation and structural and inflammatory endpoints ^19^, the present work provides a substantially expanded evaluation of preclinical efficacy. By integrating biochemical correction (including VLCFA-associated lipid abnormalities) in micro dissected whole retina and RPE, inflammatory modulation, dose-response analysis, preservation of newly recognized structural endpoints, and expanded functional endpoints, this study provides a more comprehensive and clinically relevant understanding of how retinal *PEX1* augmentation modifies disease progression.

A central biochemical feature of *PEX1* deficiency is the accumulation of VLCFA-derived lipids, including C26:0-LPC. Although it remains unknown whether these lipids are direct drivers of retinal pathology in ZSD, prior evidence suggests that VLCFA accumulation can trigger cellular stress and inflammatory responses in other peroxisome-related disorders ^47, 48^ as well as contribute to retinal pathology in other peroxisomal mouse models ^49–51^. Their substantial reduction following *PEX1* augmentation indicates that local peroxisomal restoration can partially correct the potentially toxic VLCFA overaccumulation. This biochemical improvement parallels the preservation of RPE integrity, photoreceptor structure, and visual function. Because lipid measurements reflect whole-retina averages, including non-transduced regions, the residual elevation relative to WT likely reflects the coverage limitations of subretinal AAV8 delivery ^24, 52^. Thus, while the correction does not fully normalize VLCFA levels, the observed biochemical and structural improvements demonstrate that partial metabolic rescue is sufficient to confer measurable therapeutic benefit.

Despite partial but significant correction of C26:0-LPC, DHA-containing plasmalogens remained reduced after treatment. This indicates that the degree of local peroxisomal correction achieved here is insufficient to normalize all lipid pathways affected by *PEX1* deficiency. Peroxisomes are required for DHA biosynthesis via β-oxidation of C24:6n-3 to C22:6n-3 (DHA), and peroxisomal disorders are associated with reduced tissue DHA levels ^53, 54^. Although the extent to which DHA deficiency contributes to retinal pathology in ZSD mouse models is unknown, reports suggest it is a key contributor to retinal pathology in the *Mfp2^-/-^*mouse model of peroxisomal β-oxidation deficiency ^55^. The persistence of DHA plasmalogen deficits following *PEX1* gene augmentation observed here is unlikely to reflect insufficient vector coverage, given that other lipid abnormalities were corrected under the same conditions. Instead, these findings raise the possibility that retinal DHA plasmalogens depend partly on systemic liver-derived supply, ^56–58^. While we cannot conclude from this study the effect of ongoing DHA deficiency on efficacy and durability of gene augmentation, the potential for combination therapy with oral DHA supplementation to further improve outcomes should be considered. This distinction between peroxisomal lipid metabolic pathways that can be corrected locally and those that may require systemic intervention or earlier treatment may also support and inform the use of additional combination therapy approaches, particularly in patients with more severe systemic involvement. Regardless, it is important to note that lipid profiles of ZSD patient retinal tissues are unknown, and may differ from those measured in blood or primary fibroblast cultures, as seen with the PEX1-G844D mouse model.

PEX1-G844D retinas exhibit increased subretinal mononuclear phagocytes and Müller gliosis, consistent with an inflammatory component of disease progression ^14, 15, 36^. *PEX1* augmentation reduced these inflammatory features, particularly at the 1.24 x 10^9^ vg dose, in parallel with improved structural and functional outcomes. In many IRDs, inflammation is considered a secondary response to photoreceptor loss, but in peroxisomal dysfunction models inflammatory activation can also arise from metabolic stress. Consistent with this, we observed significant recruitment of macrophage/microglial cells in the outer retina despite limited photoreceptor loss, suggesting that inflammatory activation may occur independently of evident degeneration ^59^. In the PEX1 G844D retina, the temporal relationship between metabolic dysfunction and inflammation suggests that subretinal macrophage recruitment reflects downstream consequences of early peroxisomal stress rather than being an initiating event. Our prior longitudinal characterization of this model demonstrated that lipid metabolic abnormalities and RPE structural changes arise before overt photoreceptor degeneration and precede the emergence of a robust inflammatory response ^15^. Although these observations support a model in which metabolic dysfunction is the primary trigger for inflammatory activation, our data do not establish causality. Future mechanistic studies, such as pharmacological or genetic modulation of microglial/macrophage activation in the PEX1 G844D background, will be required to determine whether inflammation contributes directly to disease progression or primarily reflects underlying metabolic stress. This interpretation is consistent with evidence that peroxisomal metabolism regulates innate immune signalling and macrophage activation ^60–62^ as well as with reports of inflammatory abnormalities in peroxisomal disorders ^63, 64^. Together, these findings support the view that inflammation is a modifiable contributor to retinal degeneration in ZSD. Finally, although inflammatory markers were reduced following treatment, our study does not resolve the extent to which inflammation contributes to disease progression in this metabolic context. Thus, determining whether direct modulation of inflammatory pathways could complement *PEX1* augmentation will require dedicated mechanistic studies.

In parallel with these efficacy findings, our WT data provide additional context for dose and promoter selection. In WT mice, ONL thinning at the injected pole occurred in both AAV8.PEX1– and AAV8.GFP–treated eyes at the maximal dose of 6.2×10 vg, indicating that high dose AAV8 exposure itself can induce retinal stress independent of the transgene. In PEX1-G844D mice, baseline ONL thinning is already severe at 7 months and subretinal delivery of AAV8.PEX1 at the maximal dose did not exacerbate ONL loss but instead showed a trend toward improved ONL preservation in the ventral retina, where vector exposure is lower. This suggests a balance between functional PEX1 restoration in regions exposed to moderate vector levels and local stress associated with high dose AAV8 exposure at the injection site. In terms of subretinal inflammation, in WT mice inflammatory cells were restricted to the injection site, and overall inflammatory response was mild, consistent with the expected local stress of subretinal injection. In contrast, PEX1 G844D mice exhibited diffuse subretinal inflammation across non injected, vehicle injected, and AAV-treated eyes, reflecting a disease associated inflammatory milieu rather than vector related toxicity. Taken together, these observations suggest that AAV8 vector- and injection-related factors are the predominant drivers of stress in the WT tissue at high dose. However, we cannot rule out the possible contribution of supraphysiologic PEX1 expression.

The CBh promoter was selected for its robust, clinically validated expression profile and to ensure sufficient rescue in a retina with constitutive PEX1 deficiency and peroxisomal dysfunction. Historically, cell studies have not shown adverse effects of *PEX1* overexpression using plasmids, and in the disease state we do not anticipate negative effects from restoring PEX1 in any retinal cell type. Notably, RPE PEX1 protein levels continued to rise with vector dose up to the highest dose tested, whereas neural retina levels plateaued across the three higher doses, indicating tissue specific saturation and supporting the need for dose titration. Disentangling potency, coverage, and toxicity would require dedicated dose-refinement studies with finer dose spacing around the 1/1-1/10 range, together with spatial correlation of PEX1 expression with local structural/functional outcomes. Although we did not formally quantify a therapeutic index in this study, this will be the focus of subsequent clinical trial enabling toxicology studies in larger animals with eyes more closely resembling human anatomy. Accordingly, the highest dose used here represents an upper boundary for preclinical evaluation rather than a clinical starting point. Future safety studies will prioritize titration to define the therapeutic index while maintaining sufficient retinal coverage for vision improvement. Moreover, although the genome integrity AAV8.*PEX1* was higher than expected for research grade vector with a payload this large ^65^, clinical trial enabling manufacturing refinement will further improve purity and may allow for lower dosing and thus reduced toxicity risk.

Several additional limitations should be acknowledged. First, although our findings show coordinated improvements in metabolic, structural, and inflammatory endpoints, the specific cellular pathways linking these processes and the key drivers of pathology remain to be clarified. Secondly, ffERG provides an average measure of retinal response across the entire tissue, including non-transduced regions, and thus may not be an accurate representation of functionally meaningful improvement. This is consistent with observations in ZSD patients that retain functional vision despite extinguished ERG ^11, 12^. Still, it remains promising that while scotopic ffERG was partially preserved compared to baseline in PEX1-G844D mice, photopic ffERG improved with treatment. In this context, outcomes including spatial acuity and contrast sensitivity are increasingly important, as correction in a subset of cells appears sufficient for functional vision improvement. Thirdly, while our results demonstrate that retinal disease progression can be modified at the age treated here, they do not define the optimal therapeutic window or the extent to which later stages of degeneration remain amenable to intervention. We selected a study endpoint of age 7 months, corresponding to a timepoint at which severe cellular degeneration and ffERG decline have occurred in our model. Although *PEX1* gene augmentation improved preclinical outcome measures to this age, we do not know the durability of these protective effects. Finally, we do not know the extent to which our preclinical efficacy observations in mice are translatable to the human disease. Ongoing prospective studies of retinal disease progression in ZSD patients will identify the optimal timepoint for intervention and outcome measures for future clinical trials ^66^.

Together, our results demonstrate that localized *PEX1* augmentation can modify the course of retinal degeneration in *PEX1* deficiency by correcting localized metabolic defects. Although full restoration of photoreceptor lipid composition was not achieved, the structural and functional improvements observed here highlight the therapeutic potential of targeting peroxisomal dysfunction within the retina, and may be applicable to additional peroxisomal disorders that feature vision loss ^67^. These findings support a model in which partial peroxisomal correction yields meaningful benefit and provides a foundation for developing other targeted genetic interventions, including gene editing, for retinal disease in peroxisomal disorders.

## MATERIALS AND METHODS

### Proviral plasmids and AAV production

A human codon-optimized *PEX1* sequence (synthesized by ATUM/DNA2.0) was amplified with Q5 DNA polymerase (New England Biolabs, Ipswich, MA) to include a Kozak consensus sequence upstream of the translational start site. The PCR product was digested with NotI-HF and ScaI-HF restriction enzymes (New England Biolabs, Ipswich, MA) and cloned into an AAV proviral plasmid using T4 DNA ligase (NEB). The completed proviral vector consisted of the CBh promoter cassette (chicken β-actin promoter/cytomegalovirus enhancer/hybrid intron) driving transgene expression and terminating in a synthetic polyadenylation signal. The expression cassette was flanked by canonical AAV2 inverted terminal repeats (ITRs). Oxford Nanopore long-read whole plasmid sequencing confirmed a full genome length of 5,074 base pairs (Plasmidsaurus, Arcadia, CA). AAV8 vectors (Center for Advanced Retinal and Ocular Therapeutics Research Vector Core, University of Pennsylvania) were generated using a previously described method ^68^ involving branched polyethylenimine (PEI) (Polysciences 23966, Warrington, PA)-mediated triple transfection of HEK293 cells with a plasmid containing the transgene between the ITRs of AAV2, an AAV-helper plasmid encoding Rep2 and Cap for serotype variants, and the pHGTI-Adeno1 plasmid harboring adenoviral helper genes. HEK293 cells express the adenoviral helper E1A/E1B gene (American Type Culture Collection CRL-157, Manassas, VA). Vectors were purified using a discontinuous iodixanol gradient (Sigma OptiPrep, St. Louis, MO). Encapsidated DNA was quantified by TaqMan qPCR, following denaturation of AAV particles by proteinase K; titers were calculated as vector genomes (vg) per mL.

### In vitro titer assay for individual AAV capsid variants

Capsid genes were cloned into an AAV packaging plasmid for vector production and used for small-scale vector preparations encoding firefly luciferase to determine titer. Physical particle titers were determined by TaqMan qPCR. Subsequently, AAV2/8bp variants were assayed for transduction at equal multiplicity of infection (MOI) in HEK293 cells. For large-scale viral titer, the encapsidated DNA was quantified by TaqMan qPCR, following denaturation of the AAV particles by proteinase K, and titers were calculated as vector genomes (vg) per mL. qPCR using CMV primer/probe indicated a concentration of 6.2 x 10^12^ vg/mL for AAV8.*PEX1* (1.7 x 10 ^13^ vg total production) and 1.18 x 10^13^ vg/mL for AAV8.GFP (1.18 x 10^13^ vg total yield). Oxford Nanopore long-read sequencing of whole AAV genomes determined 53% and 78% full-length sequence in correct orientation for AAV.*PEX1* and AAV8.GFP, respectively (Plasmidsaurus, Arcadia, CA).

### ARPE-19 cell line generation

*PEX1*-null ARPE-19 cells were generated using CRISPR-Cas9-mediated gene editing at The Jackson Laboratory (Bar Harbor, ME). Homozygous PEX1-p.[Gly843Asp] (G843D) ARPE-19 cells were generated using CRISPR-Cas9 mediated gene editing and homology-directed repair (HDR) with a single-stranded DNA donor template to introduce a biallelic (c. 2528G>A) substitution in the *PEX1* gene. Genotypes were confirmed by sequencing.

### Primary mouse RPE cell isolation

Primary retinal pigment epithelium (RPE) cells were isolated from 2-week-old mice. Enucleated eyes were incubated overnight in the dark at room temperature in DMEM (Wisent, Saint-Jean-Baptiste, Canada) containing 10% fetal bovine serum (FBS) (Wisent, Saint-Jean-Baptiste, Canada) followed by incubation for 45 minutes with 2 mg/mL trypsin/collagenase I at 37°C for gentle digestion. FBS was included to inhibit enzymatic digestion. RPE cells (8 eyes per genotype) were carefully removed from the choroid and retina (eyecup) and collected in 1 mL of complete medium (DMEM, 10% FBS, and 1% penicillin-streptomycin (Wisent, Saint-Jean-Baptiste, Canada). Cells were centrifuged 5 min at 3,500 rpm at 4°C and resuspended in 1 mL of complete medium. The cell suspension was plated on one well of a 6-well plate pre-coated with Attachment Factor solution (Cell Applications, San Diego, CA) for 30 min at 37°C. Cells were grown to confluence prior to plating for dose-response experiments. Penicillin-streptomycin was not used beyond the initial expansion phase.

### Cell transduction

ARPE-19 or primary mouse RPE cells were seeded in 12-well plates. For immunofluorescence (IF) studies, cells were grown on coverslips placed in each well. Once cells grew to confluence (∼0.5 x 10^6^ cells / well), AAV8.*PEX1* or AAV8.*GFP* vectors were added in 800uL of culture medium (DMEM + 10% FBS) at various doses. Cells were monitored daily for signs of toxicity and GFP expression.

### Immunoblotting

Flash-frozen cells or mouse tissues were lysed in RIPA buffer, separated on 4-12% Bis-Tris gradient gels (Invitrogen, Waltham, MA) and transferred to nitrocellulose membranes. Membranes were blocked and incubated in 5% milk with primary antibodies: rabbit anti-PEX1 (1:1000; Proteintech 13669-1-AP, Rosemont, IL), rabbit anti-PEX5 (1:2000; gift from Gabriele Dodt, University of Tübingen), rabbit anti-β-tubulin (1:17,000; Abcam 6046, Waltham, MA) followed by horseradish peroxidase (HRP)–conjugated goat anti-rabbit IgG (H+L) secondary antibody (Novus NB7160, Oakville, Canada). Signals were visualized by enhanced chemiluminescence (ECL) using an Amersham 600 platform. Immunoblotting was performed on cell lines 4 days post-transduction, and on mouse retina or RPE at 1, 2, and 6 months post-transduction.

### Peroxisome import after viral transduction

Five days post-AAV8 transduction, cells were prepared for indirect IF as previously described ^69^, mounted onto slides using ProLong^TM^ Gold antifade reagent with DAPI (Invitrogen, Waltham, MA), and imaged using a Leica DMI600 microscope equipped with DFC345FX camera and LAS X software (Richmond Hill, Canada). Primary antibodies included rabbit anti-human PTS1 (1:200), rabbit anti-human thiolase (1:200; gift from Steven Gould, Johns Hopkins University), 1:300 rabbit anti-human PEX5 (1:300; gift from Gabriele Dodt, University of Tübingen), rabbit anti-human catalase (1:200; AOXRE 24316, Burlingame, CA) and mouse anti-human ABCD3 (1:150; Sigma-Aldrich SAB4200181, St. Louis, MO). Secondary antibodies included donkey anti-rabbit Alexa Fluor 488 (1:400; Invitrogen A21206, Waltham, MA) and goat anti-mouse Alexa Fluor 594 (1:300; Invitrogen A11005, Waltham, MA).

### Lipid analysis

#### Cultured cells

Cells were trypsinized, centrifuged, washed twice in PBS, and flash-frozen. Cell pellets were homogenized in PBS. An extraction solution of methanol containing 10 ng each of the internal standards 16:0-D4 lyso-PAF (20.6 pmol) and 26:0-D4 lyso-PC (15.6 pmol) was added to 50 µg of protein extract in a glass tube. The samples were incubated on a shaker at room temperature for 1 hour. Samples were transferred to Corning Costar Spin-X centrifuge tube filters and centrifuged for 5 minutes. Filtrates were transferred to autosampler Verex vials (Phenomenex) for LC-MS/MS analysis.

#### Retinas

Neural retina or retinal pigment epithelia (RPE) were isolated, flash-frozen, and stored at -80°C. For biochemical analysis of peroxisomal metabolites by liquid chromatography-tandem mass spectrometry (LC-MS/MS), tissues were homogenized in PBS using a mini pestle. A 2:1 chloroform:methanol solution containing 0.05% butylhydroxytoluene (BHT) was added to 50 µg of protein extract in a glass tube and incubated on an orbital shaker at room temperature for 2 hrs. Samples were centrifuged at 2,500 rpm for 10 min, and the supernatant was transferred to a clean glass tube. The supernatant was washed with 0.2 volumes of purified water, mixed, and centrifuged at 2,000 rpm at room temperature for 5 min to separate the two phases. The upper phase was removed, and the lower phase was washed with Folch theoretical upper phase (3:48:47 chloroform:methanol:water). Samples were mixed and centrifuged at 2000 rpm for 5 min, and the upper phase was removed. The lower phase was dried under nitrogen and then in a vacuum desiccator for 30 min. The dried lipid was dissolved in 3:2 hexane:isopropanol containing 10 ng each of the internal standards, 16:0-D4 lyso-PAF (20.6 pmol) and D4-26:0 lyso-PC (15.6 pmol). Samples were filtered by centrifugation using Costar Spin-X centrifuge tube filters (Corning, Tewksbury, MA) for 5 min. Filtrates were transferred to Verex autosampler vials (Phenomenex, Torrance, CA) for analysis.

A Waters TQD triple quadrupole mass spectrometer interfaced with an Acquity UPLC (ultra-performance liquid chromatography) system was used for positive-ion electrospray (ESI)-MS/MS ionization. Chromatographic separation was achieved using a 2.1 x 50 mm, 1.7 µm Waters Acquity UPLC BEH column. Mobile phase A consisted of 54.5% water, 45% acetonitrile, and 0.5% formic acid; mobile phase B consisted of 99.5% acetonitrile and 0.5% formic acid. Both phases contained 2 mM ammonium acetate. Extracts were injected under initial solvent conditions of 85% mobile phase and 15% mobile phase B. The gradient progressed from 15% to 100% mobile phase B over 2.5 min, was held at 100% mobile phase B for 1.5 min before reconditioning the column and then returned to 85% mobile phase and 15% mobile phase B for 1 min at a solvent rate of 0.7 ml/min. A column temperature of 35°C was maintained, and injection volumes of 5uL for plasmalogen and 10 µL for lysophosphatidylcholine (lyso-PC) analyses were used. Ethanolamine plasmalogen species were detected by multiple reaction monitoring (MRM) of [M+H]+ ions with fragment ions at m/z 311, 339, 361, 385, 389, 390 corresponding to species containing 16:1, 18:1, 20:4. 22:6 and 22:4 and 18:0 at the sn-2 position, respectively. Lyso-PC species were quantified by MRM transitions corresponding to fragmentation of [M+H]+ ions to m/z 104. Authentic plasmalogen standards and the tetradeuterated internal standard 26:0-D4 lyso-PC were purchased from Avanti Polar Lipids. The tetradeuterated internal standard 16:0-D4 lyso-PAF was purchased from Cayman Chemical. HPLC-grade solvents (methanol, acetonitrile, chloroform, and water) were purchased from Fisher Scientific. Formic acid was purchased from Sigma-Aldrich. PBS was purchased from Thermo Fisher Scientific.

### Mouse husbandry

PEX1-G844D heterozygous mice were maintained on congenic 129/SvEv (129S6.Cg-Pex1^tm1.1Sjms^/Mmjax) and C57BL/6N (B6.Cg-^Pex1tm1.1Sjms^/Mmjax) backgrounds. As >80% of PEX1-G844D homozygotes die before weaning on either congenic background ^17, 18^ all experiments were performed using mice on a stable mixed background (50% C57Bl/6 and 50% 129Sv/Ev). The C57Bl/6 strain used was negative for the Rd8 mutation of the *Crb1* gene. The Rd8 mutation was removed by crossing with the Crb1^cor^ C57BL/6N strain (JAX 022521). To minimize genetic variation, we used F1 progeny generated from congenic PEX1-G844D heterozygous parents (♂C57Bl/6 x ♀129Sv/Ev or ♀C57Bl/6 x ♂129Sv/Ev). No effects of parental origin or sex were observed. Nearly all (>90%) PEX1-G844D homozygous F1 progeny survived past weaning, after which no effect of the mutation on life expectancy was observed.

Mice were housed at the RI-MUHC Glen site animal care facility with *ad libitum* access to food and water. All experiments were performed at the RI-MUHC Glen site and approved by the Research Institute of the McGill University Health Centre Animal Care Committee. Euthanasia was performed using CO_2_ under isoflurane anaesthesia (5% isoflurane in oxygen until loss of consciousness, immediately followed by CO_2_ at maximum flow rate, 4 L/min). Both male and female mice were used for all experiments, and wild-type and PEX1-G844D heterozygous mice were used as littermate controls. No phenotypic differences were observed based on sex or control genotype. Genotyping was performed as previously described ^14^.

### In vivo vector delivery

Virus was diluted to the desired dose in Pluronic F-127 buffer (Sigma-Aldrich, St. Louis, MO). Mice received meloxicam oral analgesic and were anaesthetized by intraperitoneal injection of ketamine (130mg) and xylazine (13mg) in sterile PBS per kg body weight. Proparacaine hydrochloride (Alcaine^®^, Alcon, Mississauga, Canada) was applied to the eye and pupils were dilated with tropicamide (Mydriacyl^®^, Alcon, Mississauga, Canada). Bilateral subretinal injections delivering 1μL of virus dilution per eye were performed via a trans-scleral route. A small sclerotomy was created near the ora serrata using a 30G needle. A blunt-ended 33G Hamilton syringe was inserted through the opening into the subretinal space, and virus was delivered between the RPE and neural retina. Gentamicin (0.3%) and prednisolone acetate (0.6%) ointment (PRED-G^®^, Allergan, Irvine, CA) was applied to the eyes for 2 days following surgery. Injections were performed in the animal surgical suite of the RI-MUHC Glen animal facility under a dissecting microscope. Mice were monitored for signs of discomfort or corneal injury.

### Electrophysiology

Retinal function was assessed using the Celeris^TM^ full-field flash electroretinography (ffERG) (Diagnosys, Lowell, MA). Following a 12-hr dark adaptation period, mice were anesthetized [intraperitoneal injection of ketamine (130mg) and xylazine (13mg) in sterile PBS per kg body weight]. Pupils were dilated with 1% mydriacyl tropicamide (Mydriacyl^®^, Alcon, Mississauga, CA) and anaesthetized using proparacaine hydrochloride (Alcaine®, Alcon, Mississauga, Canada). All procedures were performed in a dark room under dim red-light illumination. Recordings were performed to assess rod and cone function using scotopic (dark-adapted) ffERG and photopic (light –adapted) ffERG, respectively. All stimuli were delivered by white light (6500K (Kelvin)). Scotopic ffERGs were recorded from fully dark-adapted retinas in response to progressively brighter light stimuli (0.2, 2, 4, and 8 cd.s/m^2^; pulse frequency 1 Hz, interstimulus interval 20 s, 5 sweeps averaged per intensity, sampling frequency 2,000 Hz]. Photopic ffERGs were recorded in response to light pulses at 4, 8, and 16 cd/m^2^ [background illumination 30 cd/m^2^; pulse frequency 1 Hz, initial sweep interval 6 s, inter-stimulus interval 1 s; 10 sweeps averaged per intensity; sample frequency 2,000 Hz]. The photopic recordings were obtained 10 min after background illumination. For scotopic ffERG, a-wave amplitude was measured from baseline to the most negative trough, while b-wave amplitude was measured from the trough of the a-wave to the fourth oscillatory potential (or similar latency). For photopic ffERG, b-wave amplitude was measured from baseline to the most positive peak, consistent with the minimal a-wave in murine photopic ERGs. The c-wave amplitude was determined as the difference between baseline and the first positive peak following the a-wave.

### Functional vision testing

Functional vision in mice was assessed by measuring the optomotor reflex using a virtual-reality optomotor system (OptoDrum, Striatech, Tübingen, Germany). Freely moving mice were placed on an elevated platform and exposed to vertically oriented sine wave gratings rotating at 12 degrees/sec. When the stimulus was perceived, mice ceased body movement and tracked the grating with reflexive head movements in concert the rotation. This approach yields independent measures of right- and left-eye acuity, as only motion in the temporal-to-nasal direction evokes a tracking response. Mice were tested within 3 hrs of light onset.

To determine the <u>spatial frequency threshold</u> (visual acuity), the spatial frequency of the grating at full contrast (100%) was gradually increased from 0.011 to 0.528 cycles/degree in varying incremental steps until mice no longer exhibited tracking behavior. The highest spatial frequency eliciting a tracking response defined visual acuity for each eye (cycles/ degree).

To determine <u>contrast sensitivity</u> (lowest detectable contrast), gratings at a spatial frequency of 0.056 cycles/degree were presented while contrast was gradually reduced from 100% to 1.72% at varying incremental steps. The lowest contrast eliciting a tracking response defined contrast sensitivity for each eye (%).

Spatial frequency and contrast sensitivity were also assessed under scotopic conditions. Four filter foils were placed over each monitor to achieve 4.8 log units of light attenuation (ScotopicKit Striatech, Tübingen, Germany). Mice were tested following overnight dark adaptation, and movements recorded using an infrared camera.

### Retinal flatmount preparation and immunofluorescence

Mouse eyes were enucleated and fixed in 4% formalin for 5 min at room temperature, after which a perforation was made in the cornea with a 27G needle, and fixation continued for an additional 25 min. Fixed eyes were sectioned at the limbus, and anterior segments were discarded. Posterior eyecups consisting of the neural retina/RPE/choroid/sclera complex were collected, and the neural retina was carefully separated from the RPE/choroid/sclera complex for independent preparation. Flatmounts were incubated in PBS containing 0.1% Triton X-100 and 10% FBS (saturation buffer) for 45 min. Specimens were incubated overnight at 4°C with primary antibodies diluted in saturation buffer. Secondary antibody incubation was performed at 1hr at room temperature. Primary antibodies included rabbit anti-human cone arrestin (1:500; Millipore AB15282, Burlington, MA), rat anti-human F4/80 (1:400; Abcam ab6640, Waltham, MA), rabbit anti-human IBA1 (1:400; Wako 019-19741, Richmond, VA). Secondary antibodies included anti-rabbit Alexa Fluor 594 (1:450; Invitrogen A21207, Waltham, MA), anti-rat Alexa Fluor 647 (1:450; Invitrogen A48272TR, Waltham, MA), and anti-rabbit Alexa Fluor 488 (1:450; Invitrogen A21206, Waltham, MA). RPE flatmounts were incubated overnight at 4°C with 1:400; TRITC phalloidin (ECM Biosciences PF7551 Versailles, KY) to label the actin cytoskeleton. Neural retina flatmounts were stained with fluorescein-conjugated peanut agglutinin (1:500; Vector Laboratories, Burlingame, CA). Immunofluorescence imaging was performed using a Zeiss LSM780 laser-scanning confocal microscope. RPE cells and IBA1-positive cells were quantified in eight fields of view spanning the four poles of each RPE flatmount, excluding the central region containing the optic nerve head. Cell counts were expressed as cells/mm² to assess RPE degeneration and inflammation, respectively.

### Retinal immunohistochemistry

Eyecups from PBS-perfused mice were fixed 3 hr in 10% formaldehyde, cryoprotected in 10% (30 min on ice), 20% (1 hr on ice), and 30% (4°C overnight) sucrose in 0.1M phosphate buffer (PB), and then embedded and frozen in frozen section compound (VWR, Mississauga, Canada). Retinal cryo-sections (5µm) were blocked (1% normal goat serum, 0.1% Triton X-100, 10% FBS in PBS) for 1 hr, washed, and incubated at 4°C overnight with primary antibody diluted in incubation buffer (0.1% Triton X-100, 10% BSA in PBS). Sections were then washed and incubated with secondary antibodies for 90 min at room temperature, followed by additional washes. Coverslips were mounted using ProLong^TM^ Gold antifade reagent with DAPI (Invitrogen, Burlington, CA) and images were acquired using a Leica DMI600 microscope with a DFC345FX camera and LAS X software (Richmond Hill, Canada). Primary antibodies included rabbit anti-human PEX1 (1:400; Proteintech 13669-1-AP, Rosemont, IL), and rabbit anti-human GFAP (1:400; Abcam ab7260, Waltham, MA). Secondary antibodies were as described above. Cone photoreceptors were stained with fluorescein-conjugated peanut agglutinin (1:500; Vector Laboratories, Burlingame, CA). Central retinal cryosections (including the optic nerve head) from each eye were used to measure photoreceptor layer thickness. Twelve measurements were obtained per section at defined distances from the optic nerve head. Layer thickness was quantified using ImageJ (National Institutes of Health; http://imagej.nih.gov/). Area under the curve was calculated using Prism software (version 5.01; GraphPad Software).

### Statistical analysis

Normality was assessed for each dataset using the Shapiro-Wilk test, which provides the most reliable performance for small sample sizes. When all groups satisfied normality and exhibited comparable variances, one way ANOVA was applied. For dose-response and rescue experiments, post hoc comparisons were restricted to the untreated PEX1-G844D group (Untx) using Dunnett’s correction, which is specifically designed for multiple comparisons versus a single reference group. When at least one group failed normality or when sample sizes were too small to reliably assess distributional assumptions (n ≤ 3), nonparametric analyses were performed using Kruskal-Wallis with Dunn’s multiple comparisons test. Lipidomic datasets were analyzed using ordinary two-way ANOVA with genotype and treatment as independent factors. Datasets exhibiting a dose-response pattern were analyzed using the Jonckheere-Terpstra trend test for ordered dose groups, and results presented in **Table S1**. Longitudinal datasets (ffERG and OMR follow up) were analyzed at each timepoint. Baseline genotype differences were evaluated by comparing WT untreated to PEX1-G844D untreated. For therapeutic efficacy, all PEX1-G844D treatment groups (vehicle, AAV8.GFP, AAV8.*PEX1* at each dose) were compared to the untreated group. For injection, transgene, or capsid effects, AAV8.GFP and AAV8.*PEX1* groups were compared to vehicle treated mice of the same genotype. Statistical significance was defined as P ≤ 0.05. Mouse cohort sample sizes were based on effect sizes observed in our previous proof-of-concept study, and subsequent outcome measure refinement in place of formal *a priori* power calculations. For non-parametric analyses, effect size was reported as the Hodges-Lehmann (HL) estimator with corresponding 95% confidence intervals. For ANOVA-based analyses, effect size was reported as the predicted least-squares mean difference (LS mean difference) with corresponding 95% confidence intervals. Post-hoc power estimates were calculated for all reported pairwise comparisons. Effect sizes, confidence intervals, and post-hoc power estimates are reported in **Tables S2 - S7**. Statistical analyses were performed using GraphPad Prism 10.0 (San Diego, CA) and R version 4.6.1 (Vienna, Austria).

## Data availability statement

Numerical data including raw LC MS/MS data, processed peak tables, ffERG waveform quantification, and OMR thresholds are available in the public repository Mendeley, cited as: Omri, Samy; Argyriou, Catherine (2026), “Clinically relevant AAV8-PEX1 gene therapy preserves retinal integrity and function long-term in a murine model of Zellweger spectrum disorder”, Mendeley Data, V1, doi: 10.17632/497wgxtxw7.1. Retinal tissue images used for structural phenotyping are available at the RI-MUHC. Original immunoblot and cell images are located at the RI-MUHC and CHLA-TSRI.

## Acknowledgments

We thank the Global Foundation for Peroxisomal Disorders (GFPD) and the Wynne Mateffy Research Foundation (WMRF) for early support that made this work possible. The spirit and courage of people affected by ZSD and their loved ones are our greatest motivator. We acknowledge Costas Karatzas and the RI-MUHC technology transfer office for their guidance in securing financial support for this initiative. We thank Patricia Laplante, Ines Holzbaur, Kevin McBride, and Elizabeth Douville at AmorChem Therapeutics for their support and their critical role developing and executing a maturation plan for therapy development. We thank Ji Yun (Jenny) Song for consultation on experimental design. This work was funded by AmorChem Therapeutics (to N.E.B., C.A., J.G.H., and J.B.), the Canadian Institutes of Health Research (to N.E.B. and C.A.; CIHR 427301), the Richard and Edith Strauss Foundation (to N.E.B. and C.A.), Mitacs (to C.A.; Elevate Award IT18108), and the National Institutes of Health (to J.G.H. and N.E.B.; NIH R24 OD030033).

## Author contributions

S.O. and C.A. designed and conducted the experiments, performed statistical analyses, assembled the figures, acquired funding, and wrote the manuscript. E.D.P. performed LC-MS/MS analyses. D.S.M. designed and cloned AAV constructs. S.Z. contributed to genome integrity analyses. J.B. contributed to experimental design and provided AAV vectors. J.G.H. contributed to experimental design and assisted with manuscript writing. N.E.B. contributed to experimental design and funding acquisition. All authors contributed to manuscript review and editing.

