## Supplemental Files for "Clinically relevant AAV8-*PEX1* gene therapy preserves retinal integrity and function long-term in a murine model of Zellweger spectrum disorder"

**Table S1. Jonckheere-Terpstra (JT) Trend Analysis for Ordered Dose Groups**

| Figure | Endpoint | Groups (ordered) | JT statistic | p-value |
|---|---|---|---|---|
| **1C** | PEX1 protein expression (PEX1-null ARPE-19) | Untx → $2\times10^3$ → $10^4$ → $2\times10^4$ → $10^5$ | 74.5 | 0.0028 |
| **1C** | PEX5 protein expression (PEX1-null ARPE-19) | Untx → $2\times10^3$ → $10^4$ → $2\times10^4$ → $10^5$ | 59 | 0.1748 |
| **1D** | % Peroxisomal PTS1 (PEX1-null ARPE-19) | Untx → $2\times10^3$ → $10^4$ → $2\times10^4$ → $10^5$ | 90 | $2\times10^{-4}$ * |
| **1E** | % Cytosolic PEX5 (PEX1-null ARPE-19) | Untx → $2\times10^3$ → $10^4$ → $2\times10^4$ → $10^5$ | 90 | $2\times10^{-4}$ * |
| **1F** | % Peroxisomal catalase (PEX1-G843D ARPE-19) | Untx → $10^4$ → $2\times10^4$ → $10^5$ → $2\times10^5$ → $4\times10^5$ | 125.5 | $2\times10^{-4}$ * |
| **1G** | C26:0-LPC (PEX1-null ARPE-19) | Untx → $2\times10^3$ → $2\times10^4$ → $10^5$ → $2\times10^5$ → $4\times10^5$ | 16 | $1.727\times10^{-5}$ |
| **1G** | Total PE plasmalogens (PEX1-null ARPE-19) | Untx → $2\times10^3$ → $2\times10^4$ → $10^5$ → $2\times10^5$ → $4\times10^5$ | 119 | $1.727\times10^{-5}$ |
| **1H** | Total PE plasmalogens (PEX1-G843D) | Untx → $2\times10^3$ → $2\times10^4$ → $10^5$ → $2\times10^5$ → $4\times10^5$ | 124 | $1.079\times10^{-6}$ |
| **2A** | % Cytosolic PEX5 (PEX1-G844D primary RPE) | Untx → $2\times10^3$ → $10^4$ → $2\times10^4$ → $10^5$ | 88 | $2\times10^{-4}$ * |
| **2B** | C26:0-LPC | Untx → $2\times10^3$ → $10^4$ → $2\times10^4$ → $10^5$ | 27.5 | 0.0784 * |
| **2E** | PEX1 protein expression (RPE) | Vehicle → 1/20 → 1/10 → 1/5 → 1/1 | 70 | $9.496\times10^{-5}$ |
| **2F** | PEX1 protein expression (neuroretina) | Vehicle → 1/20 → 1/10 → 1/5 → 1/1 | 101 | $4.17\times10^{-5}$ |
| **3A** | Full contrast spatial acuity | Untx → 1/100 → 1/10 → 1/5 → 1/1 | 410 | 0.0137 * |
| **3A** | Contrast sensitivity | Untx → 1/100 → 1/10 → 1/5 → 1/1 | 184 | 0.0008 * |
| **3B** | Photopic b-wave AUC | Untx → 1/100 → 1/10 → 1/5 → 1/1 | 660 | 0.001 * |
| **4D** | ONL AUC | Untx → 1/100 → 1/10 → 1/5 → 1/1 | 51 | 0.2921 |
| **4F** | RPE cell density | Untx → 1/100 → 1/10 → 1/5 → 1/1 | 160 | 0.0018 * |
| **4G** | Subretinal macrophages | Untx → 1/100 → 1/10 → 1/5 → 1/1 | 63 | 0.0228 * |

*Permutation p-value (10 000 permutations).

**Table S2. Statistical Analyses: Functional rescue at 2 months post-treatment (Figure 3)**

| Figure | Endpoint | Comparison | Test | p-value | HL | 95% CI | Power |
|---|---|---|---|---|---|---|---|
| 3A | Full contrast spatial acuity | PEX1-G844D Untx vs PEX1-G844D Vehicle | KW + Dunn | >0.9999 | -0.01 | [-0.06 ; +0.04] | 0.10 |
| 3A | Full contrast spatial acuity | PEX1-G844D Untx vs PEX1-G844D AAV8.PEX1 1/100 | KW + Dunn | >0.9999 | -0.03 | [-0.08 ; +0.02] | 0.25 |
| 3A | Full contrast spatial acuity | PEX1-G844D Untx vs PEX1-G844D AAV8.PEX1 1/10 | KW + Dunn | >0.9999 | -0.02 | [-0.07 ; +0.03] | 0.20 |
| 3A | Full contrast spatial acuity | PEX1-G844D Untx vs PEX1-G844D AAV8.PEX1 1/5 | KW + Dunn | >0.9999 | -0.01 | [-0.05 ; +0.03] | 0.15 |
| 3A | Full contrast spatial acuity | PEX1-G844D Untx vs PEX1-G844D AAV8.PEX1 1/1 | KW + Dunn | 0.0671 | -0.04 | [-0.01 ; +0.10] | 0.30 |
| 3A | Scotopic spatial acuity | PEX1-G844D Untx vs PEX1-G844D Vehicle | KW + Dunn | >0.9999 | +0.01 | [-0.06 ; +0.08] | 0.10 |
| 3A | Scotopic spatial acuity | PEX1-G844D Untx vs PEX1-G844D AAV8.PEX1 1/100 | KW + Dunn | >0.9999 | +0.00 | [-0.06 ; +0.06] | 0.10 |
| 3A | Scotopic spatial acuity | PEX1-G844D Untx vs PEX1-G844D AAV8.PEX1 1/10 | KW + Dunn | >0.9999 | +0.02 | [-0.04 ; +0.09] | 0.15 |
| 3A | Scotopic spatial acuity | PEX1-G844D Untx vs PEX1-G844D AAV8.PEX1 1/5 | KW + Dunn | >0.9999 | +0.00 | [-0.07 ; +0.06] | 0.08 |
| 3A | Scotopic spatial acuity | PEX1-G844D Untx vs PEX1-G844D AAV8.PEX1 1/1 | KW + Dunn | >0.9999 | +0.01 | [-0.05 ; +0.07] | 0.12 |
| 3A | Contrast sensitivity | PEX1-G844D Untx vs PEX1-G844D Vehicle | KW + Dunn | >0.9999 | -2.0 | [-8 ; +4] | 0.10 |
| 3A | Contrast sensitivity | PEX1-G844D Untx vs PEX1-G844D AAV8.PEX1 1/100 | KW + Dunn | 0.4622 | +2.0 | [-2 ; +7] | 0.18 |
| 3A | Contrast sensitivity | PEX1-G844D Untx vs PEX1-G844D AAV8.PEX1 1/10 | KW + Dunn | 0.2299 | +3.0 | [-3 ; +9] | 0.20 |
| 3A | Contrast sensitivity | PEX1-G844D Untx vs PEX1-G844D AAV8.PEX1 1/5 | KW + Dunn | 0.0631 | +4.0 | [-1 ; +9] | 0.45 |
| 3A | Contrast sensitivity | PEX1-G844D Untx vs PEX1-G844D AAV8.PEX1 1/1 | KW + Dunn | 0.0163 | +6.0 | [+2 ; +10] | 0.80 |
| 3B | AUC scotopic a-wave | PEX1-G844D Untx vs PEX1-G844D Vehicle | KW + Dunn | >0.9999 | +4.0 | [-18 ; +26] | 0.10 |
| 3B | AUC scotopic a-wave | PEX1-G844D Untx vs PEX1-G844D AAV8.PEX1 1/100 | KW + Dunn | >0.9999 | -3.0 | [-23 ; +17] | 0.12 |

| | | | | | | | |
|---|---|---|---|---|---|---|---|
| 3B | AUC scotopic a-wave | PEX1-G844D Untx vs PEX1-G844D AAV8.PEX1 1/10 | KW + Dunn | 0.4325 | -9.0 | [-32 ; +3] | 0.35 |
| 3B | AUC scotopic a-wave | PEX1-G844D Untx vs PEX1-G844D AAV8.PEX1 1/5 | KW + Dunn | >0.9999 | +2.0 | [-20 ; +24] | 0.10 |
| 3B | AUC scotopic a-wave | PEX1-G844D Untx vs PEX1-G844D AAV8.PEX1 1/1 | KW + Dunn | >0.9999 | -3.0 | [-22 ; +17] | 0.12 |
| 3B | AUC scotopic b-wave | PEX1-G844D Untx vs PEX1-G844D Vehicle | KW + Dunn | >0.9999 | +72 | [-120 ; +210] | 0.12 |
| 3B | AUC scotopic b-wave | PEX1-G844D Untx vs PEX1-G844D AAV8.PEX1 1/100 | KW + Dunn | >0.9999 | -38 | [-150 ; +70] | 0.18 |
| 3B | AUC scotopic b-wave | PEX1-G844D Untx vs PEX1-G844D AAV8.PEX1 1/10 | KW + Dunn | >0.9999 | -82 | [-260 ; +40] | 0.30 |
| 3B | AUC scotopic b-wave | PEX1-G844D Untx vs PEX1-G844D AAV8.PEX1 1/5 | KW + Dunn | 0.9032 | -55 | [-190 ; +60] | 0.22 |
| 3B | AUC scotopic b-wave | PEX1-G844D Untx vs PEX1-G844D AAV8.PEX1 1/1 | KW + Dunn | >0.9999 | -40 | [-160 ; +70] | 0.18 |
| 3B | AUC Photopic b-wave | PEX1-G844D Untx vs PEX1-G844D Vehicle | KW + Dunn | >0.9999 | -4 | [-30 ; +20] | 0.15 |
| 3B | AUC Photopic b-wave | PEX1-G844D Untx vs PEX1-G844D AAV8.PEX1 1/100 | KW + Dunn | 0.8933 | -9 | [-30 ; +10] | 0.25 |
| 3B | AUC Photopic b-wave | PEX1-G844D Untx vs PEX1-G844D AAV8.PEX1 1/10 | KW + Dunn | 0.0327 | **-21** | **[-48 ; -3]** | **0.65** |
| 3B | AUC Photopic b-wave | PEX1-G844D Untx vs PEX1-G844D AAV8.PEX1 1/5 | KW + Dunn | 0.1012 | -18 | [-45 ; +5] | 0.40 |
| 3B | AUC Photopic b-wave | PEX1-G844D Untx vs PEX1-G844D AAV8.PEX1 1/1 | KW + Dunn | 0.0286 | **-22** | **[-50 ; -2]** | **0.70** |

**Table S3. Statistical Analyses: WT tolerability at 2 months post-treatment (Figure S2)**

| Figure | Endpoint | Comparison | Test | p-value | HL | 95% CI | Power |
|---|---|---|---|---|---|---|---|
| S2A | Full contrast spatial acuity | WT Untx vs WT Vehicle | KW + Dunn | 0.8376 | +0.03 | [-0.04 ; +0.10] | 0.15 |
| S2A | Full contrast spatial acuity | WT Untx vs WT AAV8-hPEX1 1/1 | KW + Dunn | >0.9999 | 0.00 | [-0.06 ; +0.06] | 0.10 |
| S2A | Full contrast spatial acuity | WT Vehicle vs WT AAV8-hPEX1 1/1 | KW + Dunn | 0.5285 | -0.02 | [-0.08 ; +0.05] | 0.18 |
| S2A | Scotopic spatial acuity | WT Untx vs WT Vehicle | KW + Dunn | >0.9999 | +0.028 | [-0.06 ; +0.11] | 0.14 |
| S2A | Scotopic spatial acuity | WT Untx vs WT AAV8-hPEX1 1/1 | KW + Dunn | >0.9999 | 0.000 | [-0.06 ; +0.06] | 0.10 |
| S2A | Scotopic spatial acuity | WT Vehicle vs WT AAV8-hPEX1 1/1 | KW + Dunn | >0.9999 | -0.028 | [-0.10 ; +0.05] | 0.17 |
| S2A | Contrast sensitivity | WT Untx vs WT Vehicle | KW + Dunn | 0.0572 | -4.56 | [-7.0 ; -0.5] | 0.55 |
| S2A | Contrast sensitivity | WT Untx vs WT AAV8-hPEX1 1/1 | KW + Dunn | 0.3212 | -2.56 | [-5.0 ; +0.2] | 0.32 |
| S2A | Contrast sensitivity | WT Vehicle vs WT AAV8-hPEX1 1/1 | KW + Dunn | >0.9999 | +2.00 | [-1.0 ; +5.0] | 0.18 |
| S2C | AUC ONL thickness | WT Untx vs WT Vehicle | KW + Dunn | >0.9999 | +200 | [-2600 ; +2600] | 0.06 |
| S2C | AUC ONL thickness | WT Untx vs WT AAV8-hPEX1 1/1 | KW + Dunn | >0.9999 | +200 | [-2600 ; +2600] | 0.06 |
| S2C | AUC ONL thickness | WT Vehicle vs WT AAV8-hPEX1 1/1 | KW + Dunn | >0.9999 | 0 | [-2600 ; +2600] | 0.05 |
| S2E | RPE cells/mm² | WT Untx vs WT Vehicle | KW + Dunn | 0.5391 | -134 | [-350 ; +50] | 0.18 |
| S2E | RPE cells/mm² | WT Untx vs WT AAV8-hPEX1 1/1 | KW + Dunn | 0.0219 | -153 | [-420 ; +20] | 0.41 |
| S2E | RPE cells/mm² | WT Vehicle vs WT AAV8-hPEX1 1/1 | KW + Dunn | 0.5391 | -21 | [-300 ; +200] | 0.12 |

**Table S4. Statistical Analyses: Retinal structural and inflammatory outcomes at 2 months post-treatment (Figure 4)**

| Figure | Endpoint | Comparison | Test | p-value | Effect size | 95% CI | Power |
|---|---|---|---|---|---|---|---|
| 4D | ONL thickness (AUC) | PEX1-G844D Untx vs PEX1-G844D Vehicle | ANOVA + Dunnett | >0.9999 | +600 | [-6261 ; +5928] | 0.10 |
| 4D | ONL thickness (AUC) | PEX1-G844D Untx vs PEX1-G844D AAV8.PEX1 1/100 | ANOVA + Dunnett | 0.0863 | -2900 | [-11528 ; +661] | 0.35 |
| 4D | ONL thickness (AUC) | PEX1-G844D Untx vs PEX1-G844D AAV8.PEX1 1/10 | ANOVA + Dunnett | 0.2204 | -2300 | [-10328 ; +1861] | 0.25 |
| 4D | ONL thickness (AUC) | PEX1-G844D Untx vs PEX1-G844D AAV8.PEX1 1/5 | ANOVA + Dunnett | 0.0498 | -5400 | [-12195 ; -5] | 0.70 |
| 4D | ONL thickness (AUC) | PEX1-G844D Untx vs PEX1-G844D AAV8.PEX1 1/1 | ANOVA + Dunnett | 0.9582 | -1400 | [-7361 ; +4828] | 0.12 |
| 4F | RPE cells/mm² | PEX1-G844D Untx vs PEX1-G844D Vehicle | ANOVA + Dunnett | >0.9999 | +73 | [-616 ; +587] | 0.10 |
| 4F | RPE cells/mm² | PEX1-G844D Untx vs PEX1-G844D AAV8.PEX1 1/100 | ANOVA + Dunnett | 0.9984 | -350 | [-707 ; +569] | 0.12 |
| 4F | RPE cells/mm² | PEX1-G844D Untx vs PEX1-G844D AAV8.PEX1 1/10 | ANOVA + Dunnett | 0.3345 | -287 | [-978 ; +225] | 0.30 |
| 4F | RPE cells/mm² | PEX1-G844D Untx vs PEX1-G844D AAV8.PEX1 1/5 | ANOVA + Dunnett | 0.0398 | -691 | [-1226 ; -24] | 0.72 |
| 4F | RPE cells/mm² | PEX1-G844D Untx vs PEX1-G844D AAV8.PEX1 1/1 | ANOVA + Dunnett | 0.1425 | -690 | [-1153 ; +123] | 0.18 |
| 4G | Subretinal MΦ/mm² | PEX1-G844D Untx vs PEX1-G844D Vehicle | KW + Dunn | >0.9999 | +25 | [-79 ; +58] | 0.10 |
| 4G | Subretinal MΦ/mm² | PEX1-G844D Untx vs PEX1-G844D AAV8.PEX1 1/100 | KW + Dunn | >0.9999 | +40 | [-67 ; +113] | 0.15 |
| 4G | Subretinal MΦ/mm² | PEX1-G844D Untx vs PEX1-G844D AAV8.PEX1 1/10 | KW + Dunn | 0.1156 | +80 | [-24 ; +129] | 0.28 |
| 4G | Subretinal MΦ/mm² | PEX1-G844D Untx vs PEX1-G844D AAV8.PEX1 1/5 | KW + Dunn | 0.1571 | +93 | [-37 ; +166] | 0.25 |
| 4G | Subretinal MΦ/mm² | PEX1-G844D Untx vs PEX1-G844D AAV8.PEX1 1/1 | KW + Dunn | 0.1502 | +94 | [-22 ; +166] | 0.22 |

**Table S5. Statistical analyses for functional rescue at 6 months post treatment (Figure 5).**

**Table S6. Statistical analyses for structural rescue at 6 months post treatment (Figures 6 and 7)**

| Figure | Endpoint | Comparison | Test | p-value | Effect size | 95% CI | Power |
|---|---|---|---|---|---|---|---|
| 6B | ONL thickness (AUC) | PEX1-G844D Untx vs PEX1-G844D Vehicle | ANOVA + Dunnett (vs Untx) | 0.9996 | +240 | [-8800 ; +9200] | 0.15 |
| 6B | ONL thickness (AUC) | PEX1-G844D Untx vs PEX1-G844D AAV8.GFP | ANOVA + Dunnett (vs Untx) | 0.9584 | +1200 | [-8000 ; +10 000] | 0.18 |
| 6B | ONL thickness (AUC) | PEX1-G844D Untx vs PEX1-G844D AAV8.PEX1 | ANOVA + Dunnett (vs Untx) | <0.0001 | -26 300 | [-34 000 ; - 18 000] | 0.99 |
| 6B | ONL thickness (AUC) | PEX1-G844D Untx vs PEX1-G844D Vehicle | ANOVA + Tukey (all pairwise) | 0.9998 | +240 | [-8 800 ; +9 200] | 0.15 |
| 6B | ONL thickness (AUC) | PEX1-G844D Untx vs PEX1-G844D AAV8.GFP | ANOVA + Tukey (all pairwise) | 0.9768 | +1 200 | [-8 000 ; +10 000] | 0.18 |
| 6B | ONL thickness (AUC) | PEX1-G844D Untx vs PEX1-G844D AAV8.PEX1 | ANOVA + Tukey (all pairwise) | <0.0001 | -26 300 | [-34 000 ; - 18 000] | 0.99 |
| 6B | ONL thickness (AUC) | PEX1-G844D Vehicle vs PEX1-G844D AAV8.GFP | ANOVA + Tukey (all pairwise) | 0.9900 | -970 | [-10 600 ; +8 700] | 0.12 |
| 6B | ONL thickness (AUC) | PEX1-G844D Vehicle vs PEX1-G844D AAV8.PEX1 | ANOVA + Tukey (all pairwise) | <0.0001 | +26 500 | [+17 800 ; +35 200] | 0.99 |
| 6B | ONL thickness (AUC) | PEX1-G844D AAV8.GFP vs PEX1-G844D AAV8.PEX1 | ANOVA + Tukey (all pairwise) | <0.0001 | +27 500 | [+18 000 ; +36 000] | 0.99 |
| 7C | RPE cells/mm² | PEX1-G844D Untx vs PEX1-G844D Vehicle | ANOVA + Dunnett (vs Untx) | 0.9625 | +53 | [-320 ; +430] | 0.12 |
| 7C | RPE cells/mm² | PEX1-G844D Untx vs PEX1-G844D AAV8.GFP | ANOVA + Dunnett (vs Untx) | 0.9999 | +8 | [-370 ; +380] | 0.10 |
| 7C | RPE cells/mm² | PEX1-G844D Untx vs PEX1-G844D AAV8.PEX1 | ANOVA + Dunnett (vs Untx) | 0.0001 | -850 | [-1200 ; - 500] | 0.95 |
| 7C | RPE cells/mm² | PEX1-G844D Untx vs PEX1-G844D Vehicle | ANOVA + Tukey (all pairwise) | 0.9788 | +53 | [-285 ; +391] | 0.18 |
| 7C | RPE cells/mm² | PEX1-G844D Untx vs PEX1-G844D AAV8.GFP | ANOVA + Tukey (all pairwise) | >0.9999 | +8 | [-330 ; +346] | 0.12 |
| 7C | RPE cells/mm² | PEX1-G844D Untx vs PEX1-G844D AAV8.PEX1 | ANOVA + Tukey (all pairwise) | 0.0002 | -851 | [-1164 ; - 538] | 0.99 |
| 7C | RPE cells/mm² | PEX1-G844D Vehicle vs PEX1-G844D AAV8.GFP | ANOVA + Tukey (all pairwise) | 0.9889 | -53 | [-330 ; +225] | 0.15 |
| 7C | RPE cells/mm² | PEX1-G844D Vehicle vs PEX1-G844D AAV8.PEX1 | ANOVA + Tukey (all pairwise) | 0.0003 | -904 | [-1210 ; - 600] | 0.99 |

| | | | | | | | |
|---|---|---|---|---|---|---|---|
| 7C | RPE cells/mm² | PEX1-G844D AAV8.GFP vs PEX1-G844D AAV8.PEX1 | ANOVA + Tukey (all pairwise) | 0.0004 | -859 | [-1180 ; -540] | 0.99 |
| 7D | GA area (mm²) | PEX1-G844D Untx vs PEX1-G844D Vehicle | KW + Dunn (vs Untx) | >0.9999 | +0.05 | [-0.32 ; +0.42] | 0.12 |
| 7D | GA area (mm²) | PEX1-G844D Untx vs PEX1-G844D AAV8.GFP | KW + Dunn (vs Untx) | >0.9999 | +0.01 | [-0.36 ; +0.38] | 0.10 |
| 7D | GA area (mm²) | PEX1-G844D Untx vs PEX1-G844D AAV8.PEX1 | KW + Dunn (vs Untx) | 0.0326 | -0.25 | [-0.40 ; -0.10] | 0.80 |
| 7D | GA area (mm²) | PEX1-G844D Untx vs PEX1-G844D Vehicle | KW + Dunn (all pairwise) | >0.9999 | +0.083 | [-0.36 ; +0.47] | 0.12 |
| 7D | GA area (mm²) | PEX1-G844D Untx vs PEX1-G844D AAV8.GFP | KW + Dunn (all pairwise) | >0.9999 | +0.010 | [-0.41 ; +0.42] | 0.10 |
| 7D | GA area (mm²) | PEX1-G844D Untx vs PEX1-G844D AAV8.PEX1 | KW + Dunn (all pairwise) | 0.0653 | +0.249 | [-0.05 ; +0.55] | 0.32 |
| 7D | GA area (mm²) | PEX1-G844D Vehicle vs PEX1-G844D AAV8.GFP | KW + Dunn (all pairwise) | >0.9999 | -0.020 | [-0.49 ; +0.40] | 0.11 |
| 7D | GA area (mm²) | PEX1-G844D Vehicle vs PEX1-G844D AAV8.PEX1 | KW + Dunn (all pairwise) | 0.8542 | +0.090 | [-0.30 ; +0.48] | 0.25 |
| 7D | GA area (mm²) | PEX1-G844D AAV8.GFP vs PEX1-G844D AAV8.PEX1 | KW + Dunn (all pairwise) | 0.2789 | +0.063 | [-0.28 ; +0.41] | 0.22 |
| 7E | MPs/mm² | PEX1-G844D Untx vs PEX1-G844D Vehicle | ANOVA + Dunnett (vs Untx) | 0.9956 | -4 | [-57 ; +50] | 0.12 |
| 7E | MPs/mm² | PEX1-G844D Untx vs PEX1-G844D AAV8.GFP | ANOVA + Dunnett (vs Untx) | >0.9999 | -1 | [-54 ; +53] | 0.10 |
| 7E | MPs/mm² | PEX1-G844D Untx vs PEX1-G844D AAV8.PEX1 | ANOVA + Dunnett (vs Untx) | 0.0376 | +46 | [+2 ; +90] | 0.75 |
| 7E | MPs/mm² | PEX1-G844D Untx vs PEX1-G844D Vehicle | ANOVA + Tukey (all pairwise) | 0.9974 | -4.0 | [-62 ; +55] | 0.12 |
| 7E | MPs/mm² | PEX1-G844D Untx vs PEX1-G844D AAV8.GFP | ANOVA + Tukey (all pairwise) | >0.9999 | -1.0 | [-59 ; +58] | 0.10 |
| 7E | MPs/mm² | PEX1-G844D Untx vs PEX1-G844D AAV8.PEX1 | ANOVA + Tukey (all pairwise) | 0.0601 | +46.0 | [-2 ; +94] | 0.45 |
| 7E | MPs/mm² | PEX1-G844D Vehicle vs PEX1-G844D AAV8.GFP | ANOVA + Tukey (all pairwise) | 0.9992 | +3.0 | [-65 ; +71] | 0.11 |
| 7E | MPs/mm² | PEX1-G844D Vehicle vs PEX1-G844D AAV8.PEX1 | ANOVA + Tukey (all pairwise) | 0.1068 | +50.0 | [-9 ; +109] | 0.40 |
| 7E | MPs/mm² | PEX1-G844D AAV8.GFP vs PEX1-G844D AAV8.PEX1 | ANOVA + Tukey (all pairwise) | 0.1376 | +47.0 | [-12 ; +106] | 0.38 |

**Table S7. Statistical analyses for biochemical rescue at 6 months post treatment (Figures 8 and S6)**

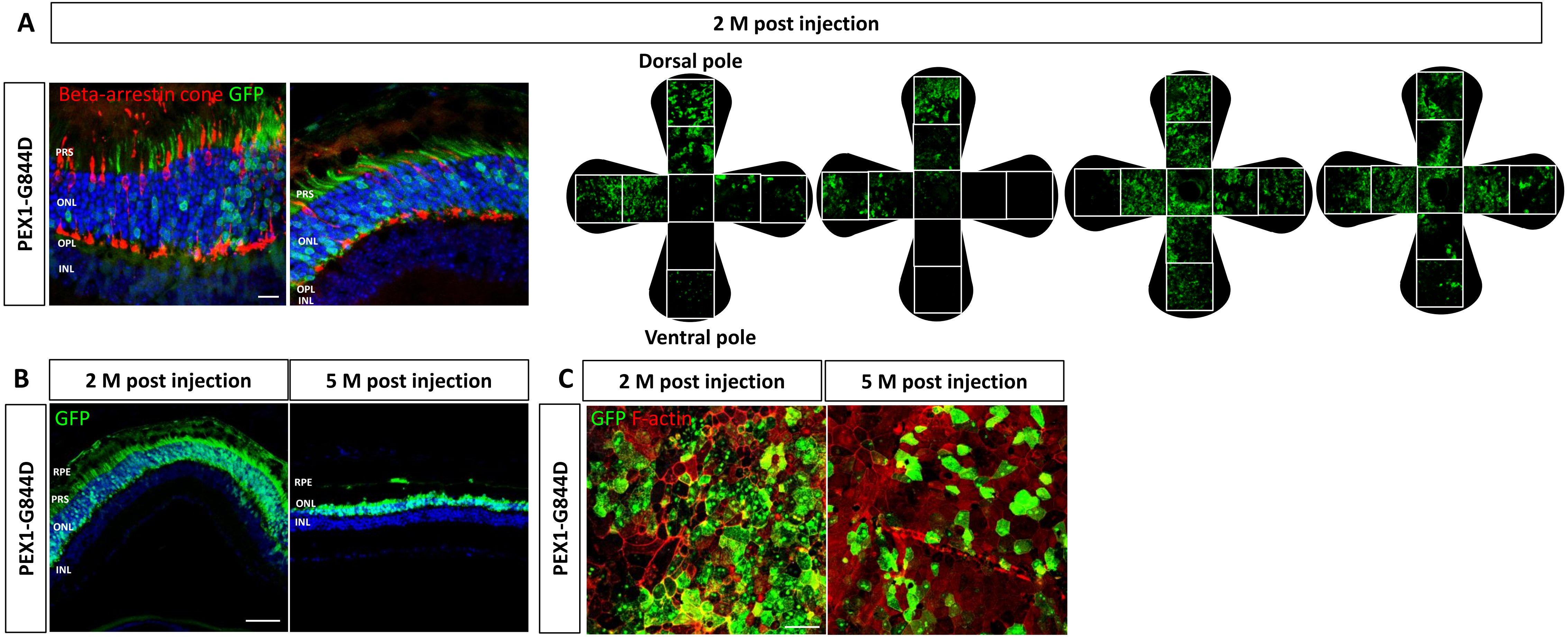


**Figure S1. Expression of AAV8-delivered GFP in PEX1-G844D mouse retina.** Representative confocal z-stack fluorescence images of **(A)** of retinal cryosections (left) or RPE flatmounts apical side up (right) 2 months post single subretinal injection of 6.2 x $10^9$vg AAV8.*GFP*, labelled with cone arrestin (red) and DAPI (blue), GFP visible in green. (scale bar: 20 µm) **(B)** Retinal cryosections (scale bar: 20 µm) and **(C)** RPE flatmounts (scale bar: 100 µm) collected 2 and 5 months after a single subretinal injection of 6.2 × $10^9$ vg AAV8.GFP, labeled with DAPI (blue) or F-actin (red); GFP shown in green. **(A,B)** PRS: Photoreceptor segments; ONL: outer nuclear layer; INL: inner nuclear layer; GCL: ganglion cell layer.

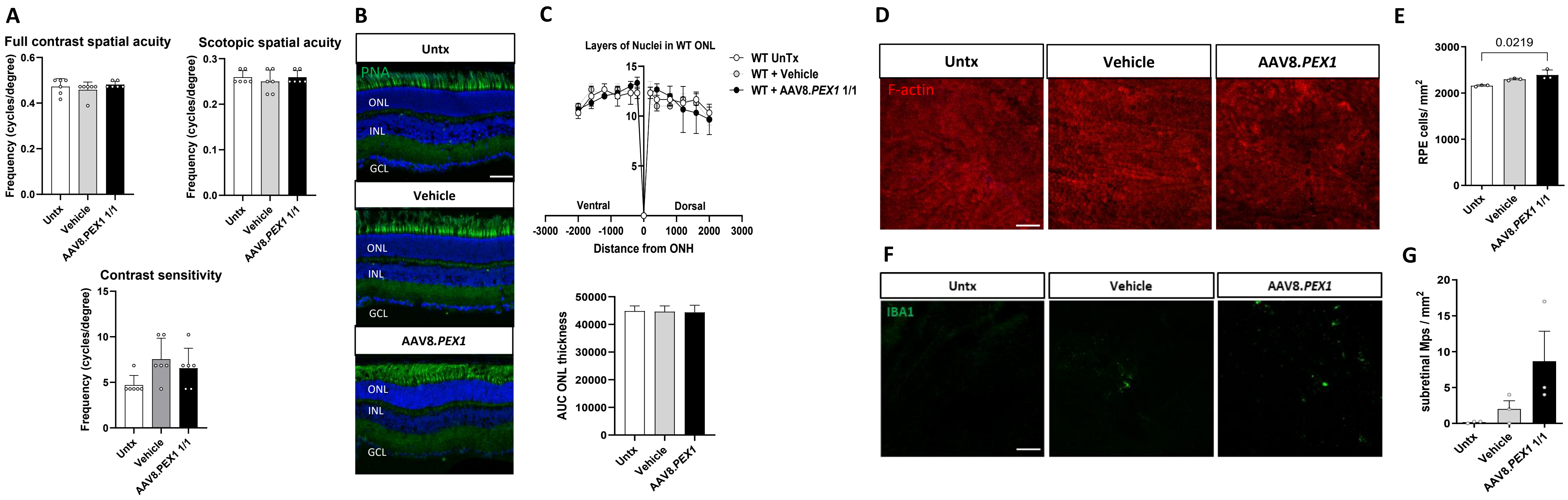


**Figure S2. Tolerability of high-dose AAV8.PEX1 in the WT mouse retina.** 6.2 x $10^9$ vg AAV8.*PEX1* or vehicle was administered by subretinal injection to 1-month-old mice, and assessments performed 2 months post treatment (age 3 months). **(A)** Optomotor reflex testing of functional vision, including full contrast spatial acuity at ambient and scotopic (low light) conditions, reported at the highest spatial frequency perceived, and contrast sensitivity at ambient light reported as the lowest contrast perceived. **(B)** Representative confocal z-stack fluorescence images (PNA in green, DAPI in blue). Scale bar = 20 µm. ONL: outer nuclear layer; INL: inner nuclear layer; GCL: ganglion cell layer. **(C)** Spider plot representation of ONL thickness across entire retinal circumference (top); Area under the curve (AUC) analysis of ONL thickness (bottom). **(D)** Representative images of RPE flatmounts (apical side up) stained with F-actin (red); Scale bar = 100 µm. **(E)** Quantification of of RPE cell density (cells/mm²). **(F)** Representative immunofluorescence images of RPE flatmounts (apical side up) labelled with IBA1 (green); Scale bar = 100 µm. **(G)** Quantification of subretinal immune cell density ($IBA1^+$ cells/mm²). N= 6 eyes for functional vision; N = 3 for histology. Kruskal-Wallis test with Dunn's multiple comparison post hoc test; P<0.05 indicated; Mean (SD) shown.

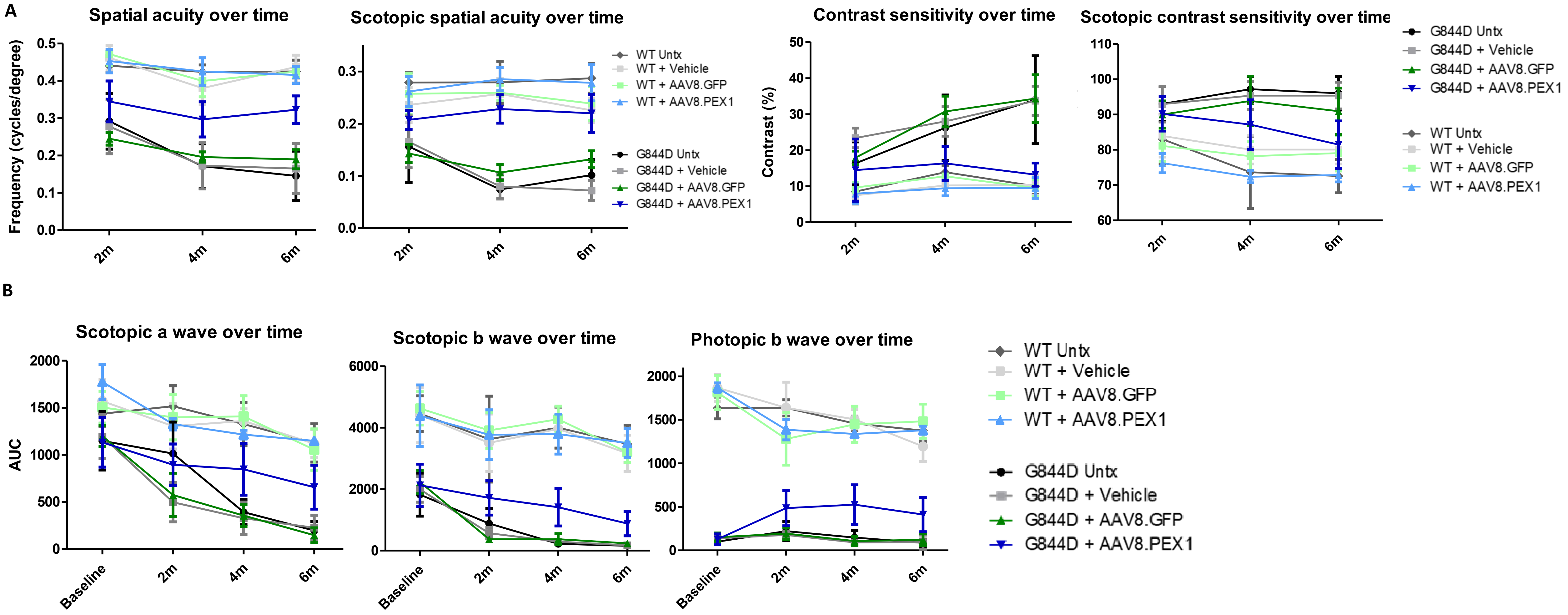


**Figure S3. Longitudinal measures of retinal function following AAV8.*PEX1* treatment in WT and PEX1-G844D mice.** Baseline ffERGs were performed on 4-week-old PEX1-G844D mice and WT littermate controls, and $1.24 \times 10^9$ vg/eye AAV8.*PEX1* or AAV8.*GFP* was administered by subretinal injection at 5 weeks of age, followed by assessments at 2, 4, and 6 months post treatment. **(A)** Optomotor reflex testing of functional vision, including full contrast spatial acuity at ambient and scotopic (low light) conditions, reported at the highest spatial frequency perceived, and contrast sensitivity at ambient light and scotopic conditions reported as the lowest contrast perceived. Values presented as a timecourse **(B)** ffERG: Area under the curve was calculated from ffERG waveforms at each timepoint and plotted as a timecourse. Data represent 6–15 mice per group (12–30 eyes). Mean (SD) shown.

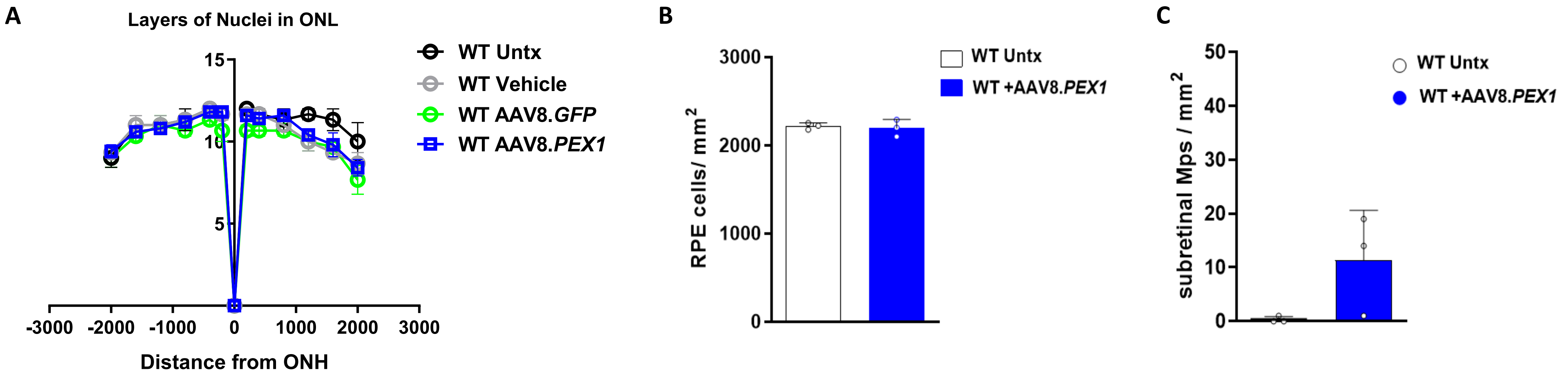


**Figure S4. Retinal integrity 6 months following AAV8.*PEX1* treatment in WT mice.** 1.24 x $10^9$ vg/eye AAV8.*PEX1* or AAV8.GFP was administered by subretinal injection to 5-week-old mice, and assessments performed 6 months post treatment (age 7 months). **(A)** Quantification of outer nuclear layer (ONL) thickness across the retinal circumference displayed as a spider plot **(B)** Quantification of RPE cell density (cells/mm²) **(C)** Quantification of the subretinal mononuclear phagocyte density (cells/mm²); N = 3 mice per group; Mann-Whitney U test; Mean (SD) shown.

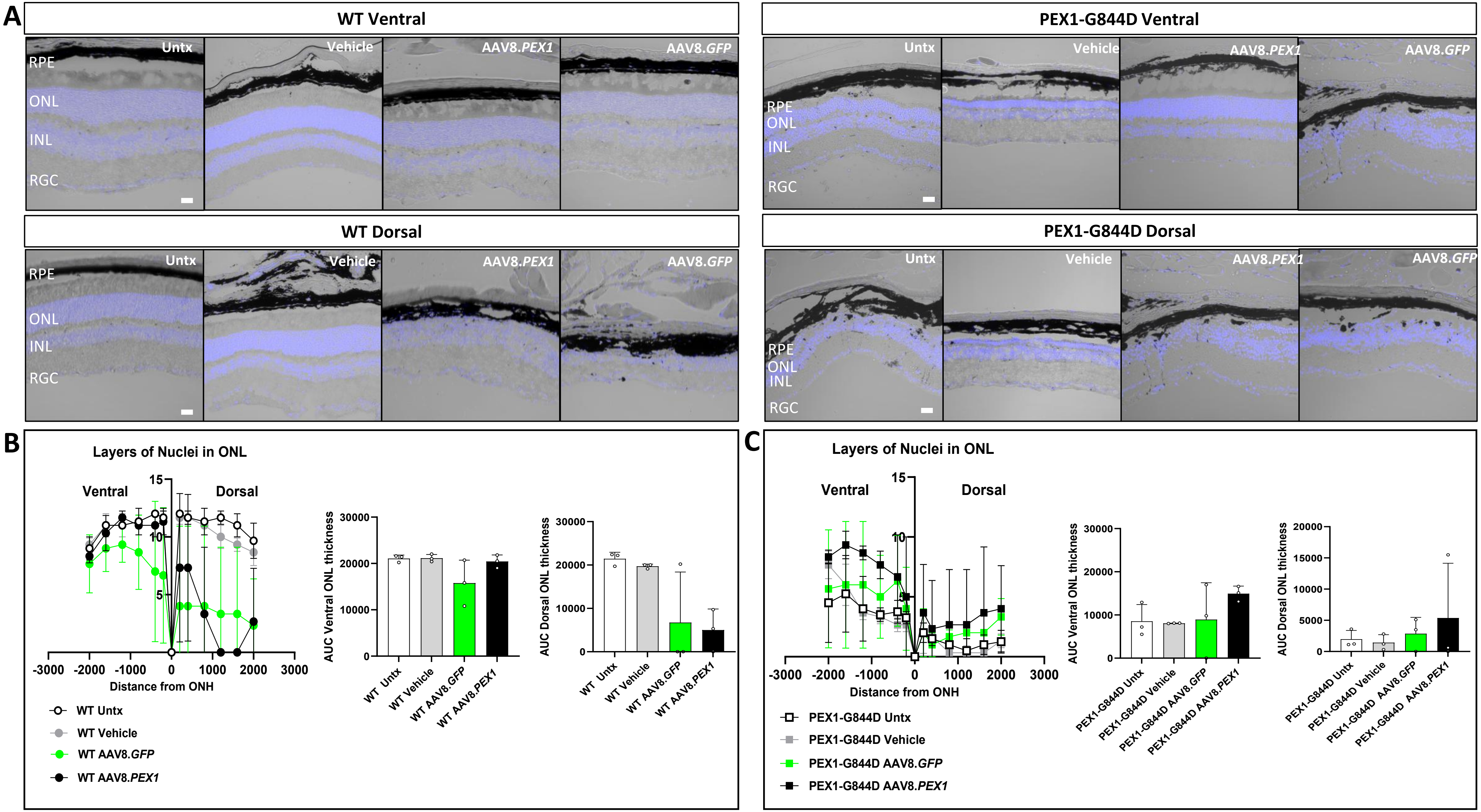


**Figure S5. Retinal integrity 6 months following high dose AAV8.*PEX1* treatment in WT and PEX1-G844D mice.** 6.2 x $10^9$ vg/eye AAV8.*PEX1*, AAV8.*GFP*, or vehicle was administered by subretinal injection to 5-week-old mice, and assessments performed 6 months post treatment (age 7 months). **(A)** Representative brightfield overlayed with confocal z-stack fluorescence images (DAPI in blue) for WT (left) and PEX1-G844D (right) ventral and dorsal retinal cryosections. Scale bar = 20 µm. RPE: retinal pigment epithelium; ONL: outer nuclear layer; INL: inner nuclear layer; GCL: ganglion cell layer. **(B)** Quantification of WT outer nuclear layer (ONL) thickness across the retinal circumference displayed as a spider plot (left); Area under the curve (AUC) analysis of ONL thickness (right). **(C)** Quantifications for PEX1-G844D retinas. N = 3 mice per group; Kruskal-Wallis test with Dunn's multiple comparison post hoc test; Mean (SD) shown.

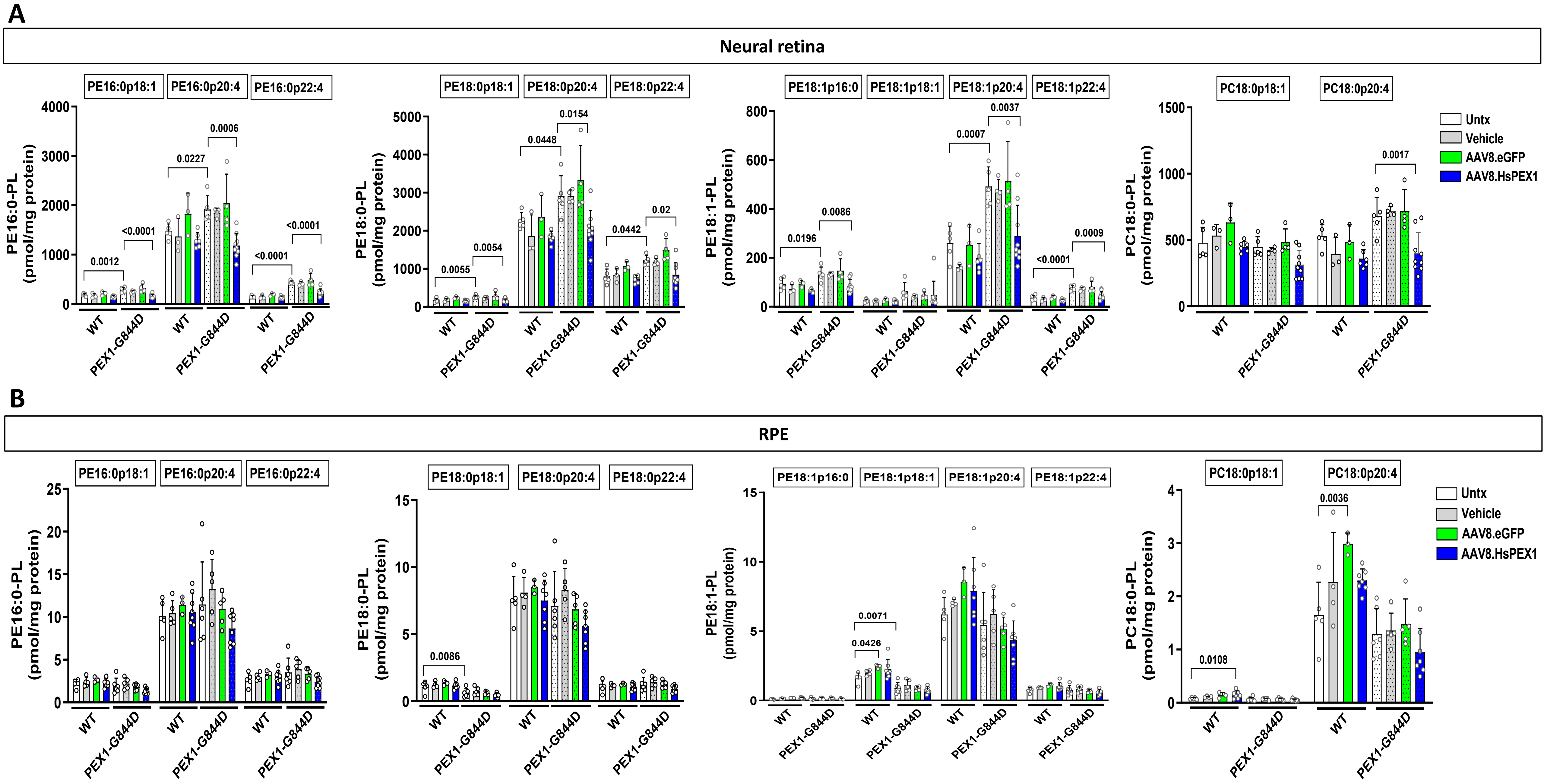


**Figure S6. LC-MS/MS quantification of peroxisomal metabolites 6 months following AAV8.*PEX1* treatment in WT and PEX1-G844D mice.** Levels of subclasses of phosphoethanolamine (PE) and phosphotidylcholine (PC) plasmalogens, were measured in **(A)** whole neural retina and **(B)** whole RPE from WT and PEX1-G844D mice 6 months following subretinal injection with 1.24 x $10^9$ vg/eye AAV8.*PEX1* or AAV8.GFP (age 7 months); N= 5-7 tissues. Two-way ANOVA with Tukey's multiple-comparisons post hoc test (cell-wise); P<0.05 indicated; Mean (SD) shown.
